# Unravelling cytosolic delivery of endosomal escape peptides with a quantitative endosomal escape assay (SLEEQ)

**DOI:** 10.1101/2020.08.20.258350

**Authors:** Serena L.Y. Teo, Joshua J. Rennick, Daniel Yuen, Hareth Al-Wassiti, Angus P.R. Johnston, Colin W. Pouton

## Abstract

Endosomal escape is an essential requirement but a major obstacle to efficient delivery of therapeutic peptides, proteins and nucleic acids. Current understanding of endosomal escape mechanisms remains limited due to significant number of conflicting reports, which are compounded by low sensitivity and indirect assays. To resolve this, we developed a highly sensitive Split Luciferase Endosomal Escape Quantification (SLEEQ) assay to probe mechanisms of cytosolic delivery. We applied SLEEQ to evaluate the endosomal escape of a range of widely studied putative endosomal escape peptides (EEPs). We demonstrated that positively-charged EEPs enhanced cytosolic delivery as a result of increased non-specific cell membrane association, rather than increased endosomal escape efficiency. These findings transform our current understanding of how EEPs increase cytosolic delivery. SLEEQ is a powerful tool that addresses fundamental questions in intracellular drug delivery and will significantly improve the way materials are engineered to increase therapeutic delivery to the cytosol.

## Introduction

Biological therapies such as peptides, proteins and nucleic acids have emerged as promising approaches for combating a wide variety of infectious, immunological and genetic disorders^1,2^. In order to elicit a therapeutic response, these macromolecules need to interact with their corresponding targets that are often located within the cell. However, unlike many low molecular weight drugs, they do not readily diffuse across cell membranes due to their large size. Instead, they are typically taken up into cells by endocytosis^3–5^. Through this pathway, most macromolecules remain trapped within membrane-bound endocytic vesicles, separating them from their required site of action. Thus, overcoming endosomal entrapment is vital for successful therapeutic delivery. However, the process of endosomal escape is inefficient and a major rate-limiting step in intracellular delivery of biological therapies^3,4^.

During the last 30 years, endosomal escape peptides (EEPs), also known as cell-penetrating peptides (CPPs), have emerged as promising delivery agents for enhancing endosomal escape. Although there are over 7,000 research papers in the growing body of literature, the mechanisms by which EEPs facilitate cytosolic delivery are yet to be fully characterised and remain the subject of significant debate^6,7^. Initial reports suggested that EEPs enter cells via direct translocation^8^ across the plasma membrane, but subsequent re-evaluation studies indicate that uptake of EEPs occurs by endocytosis^9^. Following endocytic uptake, EEPs are thought to gain entry to the cytosol by promoting endosomal membrane fusion and destabilisation^10,11^. While this has been the widely accepted explanation, a number of reports have argued against the ability of EEPs to induce endosomal release after cellular uptake^12,13^. To date, there is no clear consensus on how EEPs mediate intracellular delivery. This remains a critical issue that must be addressed in order to design more effective delivery vectors that promote endosomal release.

The uncertainty of endosomal escape mechanisms stems from the lack of methods that can quantify endosomal escape directly and reliably. Assays that measure biological activity of biomolecules are indirect approaches as they require a cascade of events to occur following endosomal escape (e.g. transcription/translation or nuclear translocation)^5,14^. In these assays, it is challenging to decouple endosomal escape from inefficiencies in these downstream processes.

Alternative methods rely on observing intracellular distribution of fluorescently-labelled materials by fluorescence microscopy. While this approach provides a direct visualisation of subcellular localisation of materials, the main disadvantage is that it is challenging to observe weak, diffuse signal—indicating cytosolic delivery—in the presence of bright, punctate signal in endo/lysosomes. Furthermore, any punctate signal that does not colocalise with common endosomal markers such as Rab5 or EEA1 (early endosome), Rab7 (late endosome) and LAMP1 (lysosome) is often mischaracterised as endosomal escape. Co-incubation of materials of interest with calcein, a small membrane-impermeable dye that appears punctate when sequestered within endosomal/lysosomal compartments, is often another method used to overcome the requirement for fluorescent labelling. However, this approach does not provide a direct measurement of cargo escape. Determining what constitutes endosomal escape is therefore highly subjective.

The major disadvantage of both of these fluorescent localisation methods is they only provide qualitative assessment of endosomal escape (yes/no) but not quantitative information (i.e. they do not measure what proportion of material taken up has escaped). It is challenging to quantify the fluorescence intensity of the cytosolic material, as both cytosolic material and that sequestered in the endosomes/lysosomes will exhibit fluorescence. To overcome this limitation, split green fluorescence protein (GFP) was developed as an endosomal escape probe that allows cytosolic signal to be distinguished from sequestered signal^15–17^. This approach provides a direct quantification of cytosolic delivery, but is limited by poor sensitivity. Typically, concentrations above 10 μM are required to observe endosomal escape, but these concentrations are significantly higher than therapeutically and clinically relevant doses for most biological materials. Due to limitations with the sensitivity of the existing assays, it is not clear if these high concentrations are required to induce escape, or if the high concentrations are simply required to generate enough signal to detect escape.

To date, there has not been a direct, highly sensitive and quantitative assay that can distinguish endosomal sequestration from cytoplasmic distribution. There is a significant need for a robust endosomal escape assay that: i) directly measures cytosolic delivery of the therapeutic; ii) is highly sensitive so it can detect the low concentrations of material delivered to the cytosol; iii) is quantitative; and iv) can determine both the amount of material that escapes and the efficiency of escape.

To address this, we developed a highly sensitive assay for the quantification of endosomal escape based on a split NanoLuciferase reporter system, termed ‘Split Luciferase Endosomal Escape Quantification’ (SLEEQ) (Fig. 1). The split luciferase assay comprises of two subunits: Large BiT protein (LgBiT, 17.8 kDa) and a high affinity complementary peptide (HiBiT, 1.3 kDa)^18^. We expressed LgBiT as a fusion protein with actin, which localises the LgBiT in the cytosol. HiBiT was attached to a protein of interest (GFP) to quantify transport to the cytosol. We demonstrated that SLEEQ can be used to detect picomolar concentrations of proteins delivered to the cytosol, and can quantify the efficiency of endosomal escape. Endosomal escape is a highly inefficient process, with only ~2% of GFP reaching the cytosol in HEK293 cells, and ~7% of GFP reaching the cytosol in HeLa cells. We also applied SLEEQ to explore the endosomal escape efficiency of a range of putative EEPs. While positively charged EEPs increased the total amount of protein delivered to the cytosol, the efficiency of endosomal escape was the same or lower than the efficiency of GFP escape without EEPs. This suggests that the positively charged EEPs increase cytosolic accumulation mostly through non-specific association with the cells, rather than inducing an active mechanism of endosomal escape. Since the EEPs studied here do not increase endosomal escape, a more appropriate name for this group of peptides would be membrane adsorptive peptides (MAPs).

**Figure 1.**
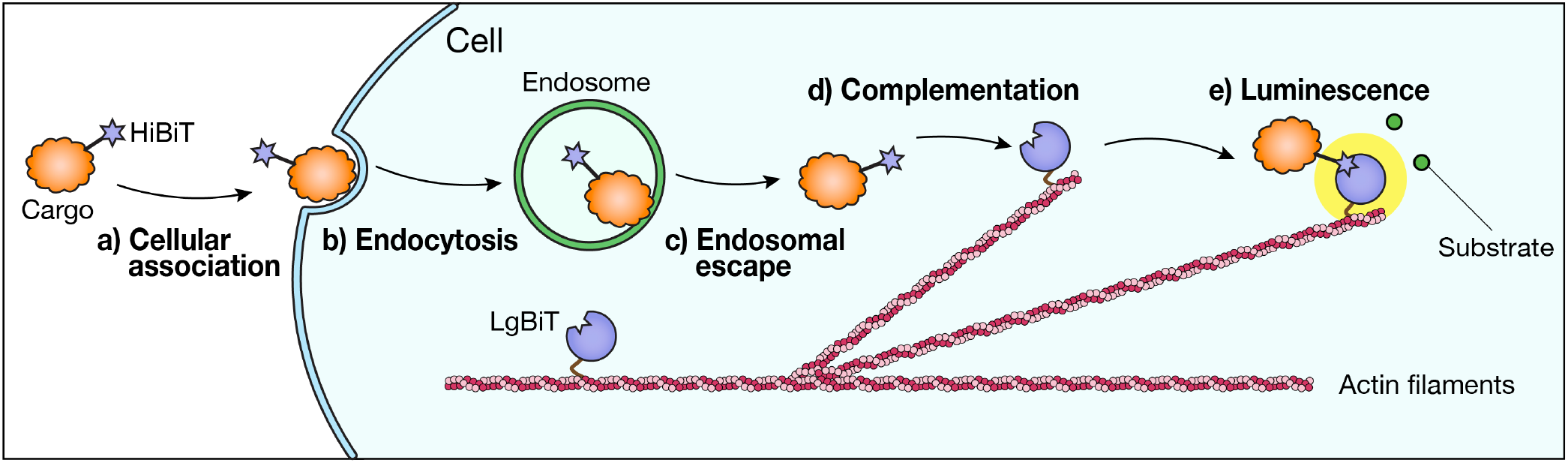
The Split Luciferase Endosomal Escape Quantification (SLEEQ) assay enables quantification of endosomal escape in live cells. a) Therapeutic cargo is labelled with HiBiT peptide. b) Endocytosis of the HiBiT-tagged cargo results in accumulation in the endosomes. c) If endosomal escape occurs, HiBiT-tagged cargo can bind to d) LgBiT protein, which is fused to actin filaments to restrict localisation to the cytosol. e) Complementation between HiBiT and LgBiT forms a functional luciferase enzyme complex, which gives off bright luminescence in the presence of a substrate.

## Results

### SLEEQ is an ultra-sensitive assay

Given that endosomal escape is a very inefficient process, it is essential to have an assay that can detect very low concentrations of cytosolic material. Split GFP systems have been employed to detect cytosolic delivery of EEPs^16,17,19^, however the sensitivity of fluorescence techniques is typically limited to micromolar concentrations. To demonstrate the sensitivity of the SLEEQ assay, we compared split NanoLuciferase to split GFP by incubating different concentrations of the small peptide fragment (HiBiT or GFP_11_) with an excess of the larger protein fragment (LgBiT or GFP_1-10_) (Fig. 2a). Our results show a linear correlation with luminescence down to 5 pM of HiBiT for the split NanoLuciferase assay, which is more than 4 orders of magnitude more sensitive than the detection limit for split GFP (limit of detection = 0.3 μM).

**Figure 2.**
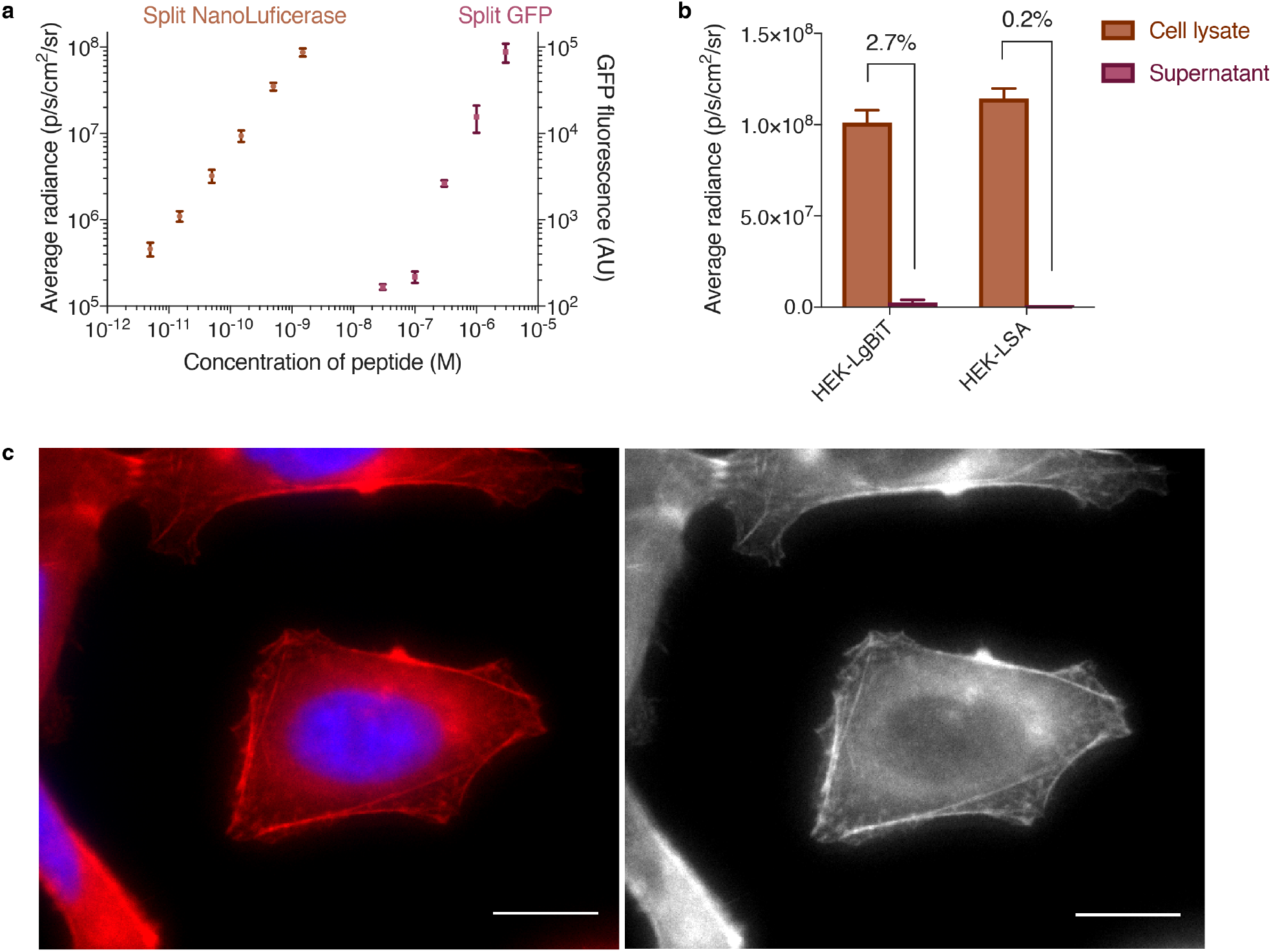
SLEEQ is four orders of magnitude more sensitive than split-GFP. a) In a 96-well plate assay, the concentration of LgBiT and GFP_1-10_ were fixed at 50 nM and 3 μM respectively. LgBiT was sensitive to < 5 pM of HiBiT, while GFP_1-10_ required > 0.1 μM of GFP_11_ to generate a detectable signal. b) HEK293 cells with LgBiT expressed in the cytosol (HEK293-LgBiT) excreted 2.7% of the protein into the supernatant. When LgBiT was fused to cytoskeletal protein actin, (HEK293-LSA) only 0.2% of the LgBiT was detected in the supernatant. A SNAP-tag was also fused to LgBiT-actin to aid visualisation of the fusion protein. c) Pseudocoloured (left) and grayscale (right) images of HEK293 cells expressing LSA (red) show cytosolic staining consistent with actin. Nucleus was stained with Hoechst 33342 (blue). Scale bar = 10 μm.

### Development and validation of SLEEQ

Having demonstrated the superior sensitivity of split NanoLuciferase, we developed SLEEQ to investigate endosomal escape in mammalian cells. First, we engineered cell lines to express LgBiT protein in the cytosol. HEK293 and HeLa cells were transduced with a lentiviral vector encoding the LgBiT transgene and expression of LgBiT was measured using luminescence by lysing the cells and adding HiBiT peptide with fumarizine substrate. Luminescent signal in transduced cells (1.01 x 10^8^ p/s/cm^2^/sr) was 732 times higher than in non-transduced cells (1.38 x 10^5^ p/s/cm^2^/sr), indicating successful expression of LgBiT protein (Supplementary Fig. 1). Low background levels of luminescence were observed in transduced cells without added HiBiT (1.04 x 10^5^ p/s/cm^2^/sr). These results demonstrate that both HiBiT peptide and LgBiT protein exhibits minimal background signal on their own. However, we detected significant levels of LgBiT protein in the cell media after 24 hours (Fig. 2b), which is likely due to secretion of LgBiT protein from the cells. Although secreted LgBiT can be washed from the cells before measuring luminescence, it is possible that the LgBiT/HiBiT complexes could form outside the cells, then be endocytosed by the cell, giving a false estimate of endosomal escape. To limit this, LgBiT was fused to β-actin, a protein found exclusively in cell cytosol. A SNAPtag was also incorporated to allow visualisation of LgBiT incorporation into actin filaments by fluorescence microscopy. Two single cell clones of HEK293 and HeLa cells were established that stably express LgBiT-SNAP-actin (LSA). Cell lysate from the LSA cell lines treated with HiBiT peptide showed comparable luminescence to the free LgBiT (Fig. 2b), suggesting HiBiT still forms an active complex with LgBiT protein when fused to actin. Secretion of LSA into the cell media over 24 hours was < 0.2% of the total expression levels in the stable cell lines, which was 10 times lower than secretion of free LgBiT from LgBiT-expressing cells (Fig. 2b), confirming LSA sequestration in the cytoplasm.

Distribution of LgBiT protein throughout the cytosol was confirmed by fluorescence microscopy. HEK293-LSA cells were treated with SNAP-Cell 647-SiR to label the SNAPtag fused to actin. Fig. 2c shows the typical pattern for actin staining, with distinct filaments throughout the structure of the cell, and no discernible punctate fluorescence. The concentration of LSA expressed in the cytosol was estimated to be 55.8 nM by permeabilising the cells with 0.01% w/v digitonin (Supplementary Fig. 2a, c and d). The presence of 0.01% w/v digitonin did not affect the LgBiT luminescence (Supplementary Fig. 2b). A number of endocytic pathways involve actin, therefore it is possible that fusing LgBiT to actin could affect protein uptake. To investigate this, we compared the fluid phase endocytosis of wild type HEK293 and HEK293-LSA cells using calcein, a small impermeable fluorescent dye. There was no significant difference in the uptake of calcine in these two cell lines, (Supplementary Fig. 3), suggesting that endocytosis is not significantly affected by the presence of the LgBiT actin fusion.

### Cationic EEPs increase cytosolic delivery of GFP

Next, we applied SLEEQ to investigate the endosomal escape of a range of putative endosomal escape peptides (EEPs) fused to green fluorescent protein (GFP) as a model delivery cargo. Each EEP has a different length and amino acid composition that therefore varies the total net charge of the proteins (Fig. 3c). We chose eight widely used EEPs that have reported endosomal escape capabilities. TAT^20,21^, polyarginine (R9)^22,23^, 5.3^24^ and ZF5.3^24^ are cationic arginine containing peptides. The 5.3 peptide incorporates five arginine residues along three helical faces of the avian pancreatic peptide scaffold whereas ZF5.3 incorporates this arginine topology into a zinc finger domain^24^. ZF5.3 has been shown to induce higher levels of cytosolic delivery than 5.3 when fused to SNAP-tag^25^. We also selected ampiphilic peptides including pHlip^26,27^, pHD118^28^ and HA2^29^. These peptides are thought to depend on acidification post-internalisation to induce endosomal escape and are commonly referred to as pH-dependent membrane-active peptides (PMAPs). pHlip is derived from bacteriorhodopsin^26,27^, whereas pHD118 is a derivitative of bee venom (melittin)^28^. The influenza haemagglutinin N-terminal peptide HA2 has also been investigated widely, with substitution of certain amino acids with glutamic acid providing improved pH dependent activity (such as E5)^29,30^. Combination of E5 and TAT (E5TAT peptide) results in a dual arginine rich and pH sensitive peptide which ideally employs both arginine driven cellular association with pH dependent membrane disruption to increase endosomal escape efficiency^31^.

**Figure 3.**
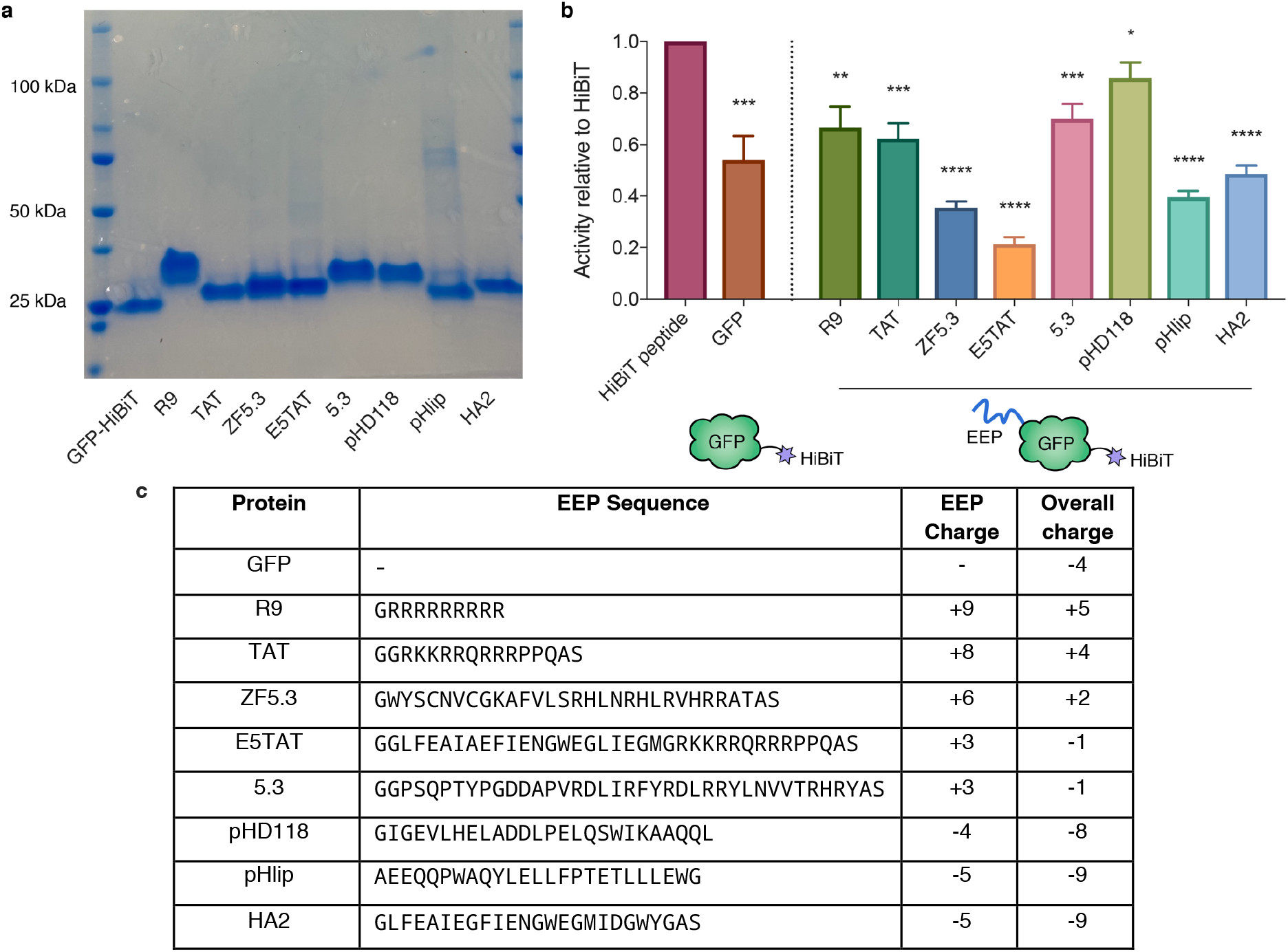
HiBiT fused to EEP modified GFP retains luminescent activity. a) SDS-PAGE gel of EEP-GFP-HiBiT fusion proteins. b) Luminescent activity of EEP-GFP-HiBiT relative to HiBiT peptide (activity of HiBiT normalised to 1). For simplicity in naming, EEP-GFP-HiBiT fusion proteins are labelled by the EEP. Data represents mean ± SEM, n = 3. * denotes p ≤ 0.05, ** denotes p ≤ 0.01, *** denotes p ≤ 0.001 and **** denotes p ≤ 0.0001. c) Summary of EEP peptide sequences, their respective charges and the overall charge when fused to GFP-HiBiT. EEPs are listed in order of decreasing charge.

The EEPs were fused to the N-terminus of GFP while HiBiT was fused at the C-terminus. SDS-PAGE shows the purity of the protein yield (Fig. 3a), and the protein size was confirmed by MALDI (Supplementary Table 2).

Fusing HiBiT to another protein may affect its luminescent activity. To control for this, the relative activity of each EEP-GFP-HiBiT fusion protein was compared to HiBiT peptide by combining the fusion proteins with an excess of purified recombinant LgBiT protein. The luminescent signal for the fusion proteins was between 20% and 80% of the free HiBiT peptide (Fig. 3b). Given the sensitivity of HiBiT peptide is ~5 pM, we determined the loss in activity would not significantly affect the sensitivity of the assay.

Next, we investigated the cytosolic accumulation of these fusion proteins in HEK293 cells. After 4 hours of incubation with 1 μM EEP-GFP-HiBiT, the cells were washed to remove unbound material and luminescence was measured.

The EEP-GFP-HiBiT proteins exhibited minimal cell toxicity at this concentration, and the cells remained viable throughout the experiment (Supplementary Fig. 4). To account for the differences in HiBiT activity for each EEP-GFP-HiBiT protein (Fig. 3b), the cytosolic and total cellular signal was normalised to the activity of free HiBiT (Fig. 4a and b). GFP without an EEP served as a baseline for endosomal escape, as any cytosolic delivery of GFP should be due to constitutive levels of cytosolic transport.

**Figure 4.**
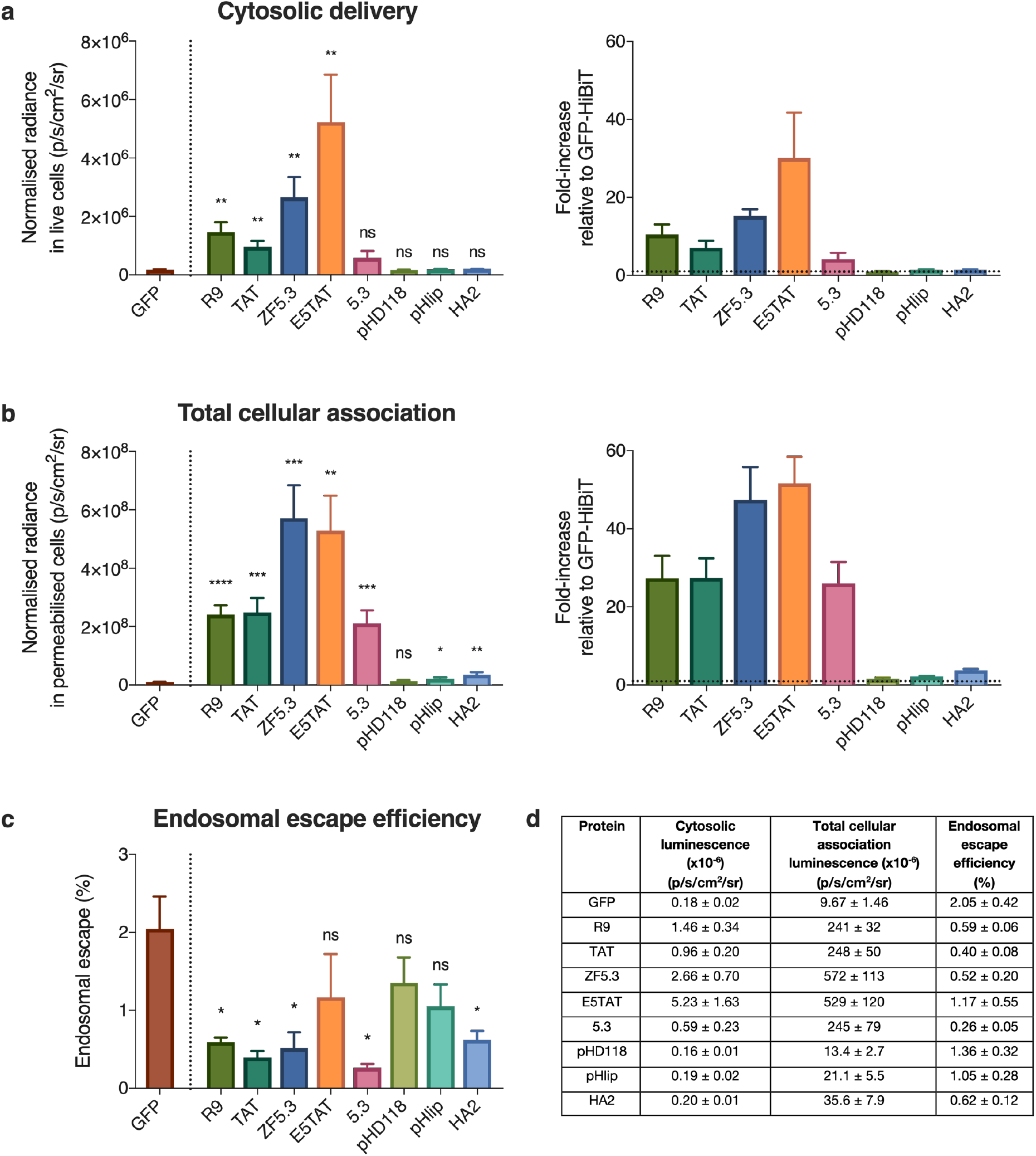
Cationic EEPs increase cytosolic delivery of GFP but do not increase endosomal escape efficiency. a) Cytosolic luminescent signal of EEP-GFP-HiBiT in HEK293-LSA cells and fold-increase in signal with respect to GFP (represented by dotted line = 1). HEK293-LSA cells were incubated with EEP-GFP-HiBiT proteins at 1 μM for 4 hours. b) Total cellular association of EEP-GFP-HiBiT in HEK293 cells and fold-increase with respect to GFP (represented by dotted line = 1) determined by permeabilising the cells using 0.01% w/v digitonin. c) Endosomal escape efficiency of EEP-GFP-HiBiT proteins determined by ratioing cytosolic signal with total cellular association. d) Summary of cytosolic luminescence, total cellular association luminescence and endosomal escape efficiency for all proteins. Data represents mean ± SEM, n=3 (except for GFP, where is n=5). Student’s t test was used to compare differences between EEP-GFP-HiBiT fusion protein and GFP. ns (not significant) denotes p > 0.05, * denotes p ≤ 0.05, ** denotes p ≤ 0.01, *** denotes p ≤ 0.001 and **** denotes p ≤ 0.0001.

R9, TAT, ZF5.3 and E5TAT showed significantly enhanced delivery (7 to 30-fold increase) of GFP to the cytosol (Fig. 4a). Surprisingly, cytosolic delivery of 5.3, pHD118, pHlip, and HA2 was not significantly higher than GFP alone (Fig. 4a). As expected, GFP showed low delivery to the cytosol. A similar trend was observed in HeLa cells (Supplementary Fig. 5a).

### Cationic EEPs enhance total cellular association

While R9, TAT, ZF5.3 and E5TAT peptides significantly increased cytosolic accumulation compared to GFP without an EEP, all these EEPs have a positive charge. It is well established that positively charged proteins associated strongly with negatively charged plasma membranes^32^. Therefore, to decouple cytosolic accumulation from the total amount of protein adsorbed to the cell, we determined the total cellular association (both surface-bound and internalised material) of the proteins. To do this, cells were washed stringently after incubation to remove all unbound materials and treated with digitonin for 1 hour to permeabilise all cellular membranes (Fig. 4b).

As expected, the positively charged EEPs (R9, TAT, ZF5.3, E5TAT and 5.3) showed significantly higher cellular association (29 to 49-fold) than GFP (Fig. 4b). In HeLa cells, a similar trend with the total cellular association was observed (30 to 79-fold, Supplementary Fig. 5b). Negatively charged EEPs (pHD118, pHlip and HA2) showed similar association to GFP in both cell lines. This highlights how positively charged EEPs can significantly influence interaction with the plasma membrane.

To further investigate the intracellular distribution of EEP-GFP-HiBiT, the cells were imaged using confocal microscopy (Fig. 5). As predicted by the cellular association results (Fig. 4b), the fluorescence signal from the negatively charged proteins was significantly lower than the signal from the positively charged proteins. To examine the distribution of EEP-GFP-HiBiT within the cells, the dynamic range of each image was optimised to aid visualisation. Unadjusted images for comparison of signal intensity can be found in Supplementary Fig. 6. All proteins showed distinct, punctate fluorescence, indicating entrapment within endosomal/lysosomal compartments and minimal endosomal escape. E5TAT also displayed pronounced membrane association on the cell surface.

**Figure 5.**
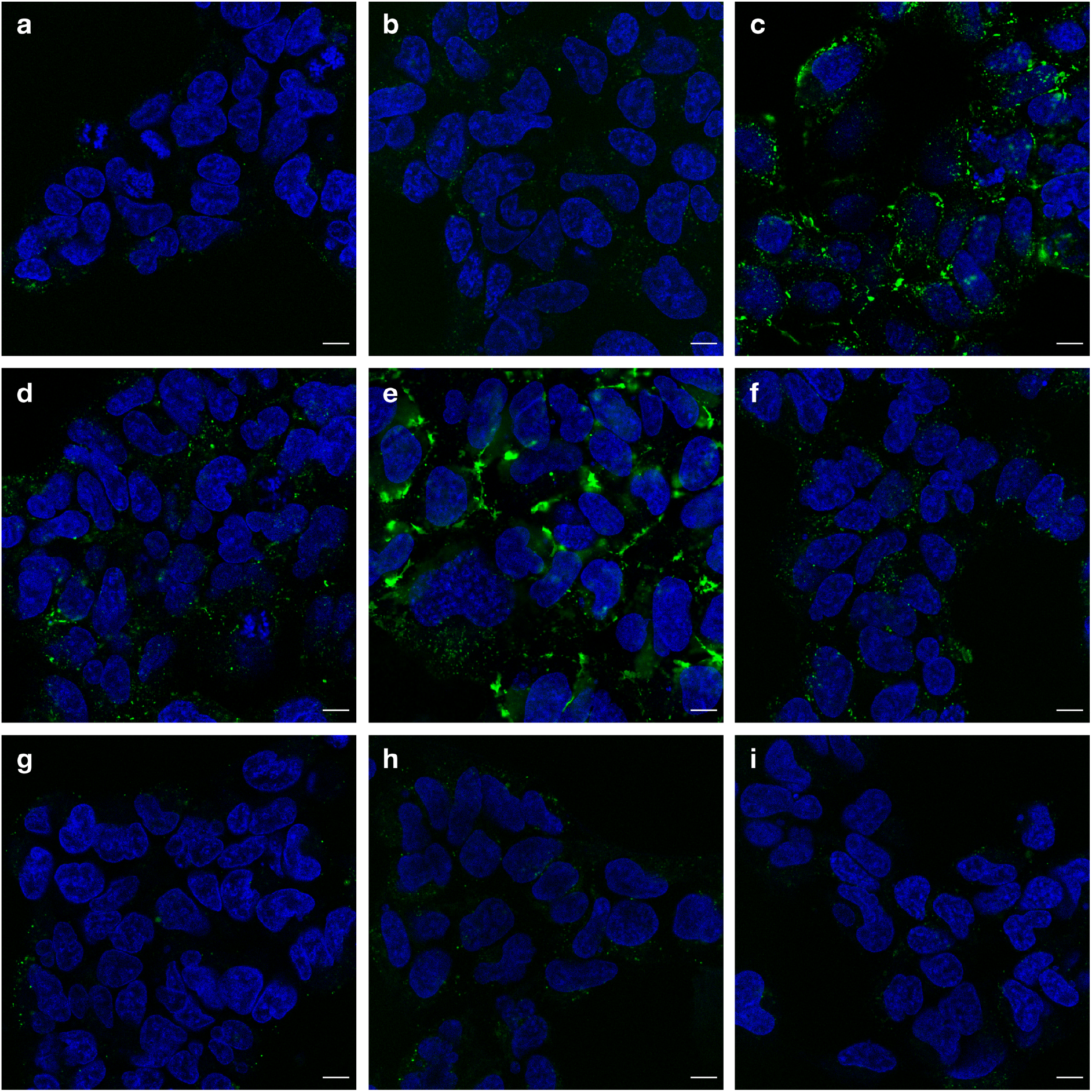
EEP-GFP-HiBiT fusion proteins exhibit punctate staining, suggesting limited endosomal escape. HEK293-LSA cells treated with 1 μM EEP-GFP-HiBiT (green) proteins: a) GFP, b) R9, c) TAT, d) ZF5.3, e) E5TAT, f) 5.3, g) pHD118, h) pHlip and i) HA2 at 1 μM for 4 hours. Cells were washed and nuclei were stained with Hoechst 33342 (pseudocoloured blue) before confocal microscopy. Scale bar = 10 μm.

To determine the endosomal escape efficiency of the different EEPs, we calculated the ratio of the cytosolic signal divided by the total associated signal (Fig. 4c). The endosomal escape efficiency of GFP alone was approximately 2% in HEK293 cells, which was lower than that of HeLa cells (~7%, Supplementary Fig. 5c and d). Strikingly, none of the EEPs tested had higher endosomal escape efficiency than GFP alone. Even more significantly, while several of the positively charged EEPs showed higher total cytosolic delivery, the endosomal escape efficiency was significantly lower than GFP alone. Overall, these results suggest that none of the EEPs tested were more efficient at delivering protein to the cytosol than the constitutive cytosolic transport mechanisms.

While low endosomal escape efficiency was observed for all EEPs, it is possible that the high cell association could be saturating the endosomal escape capacity of the cells. To probe this, we attempted to match total cellular association of the proteins by adjusting the concentrations of protein added to the cells (Fig. 6). Free HiBiT peptide was also included in these experiments to determine if HiBiT has any endosomal escape properties.

**Figure 6.**
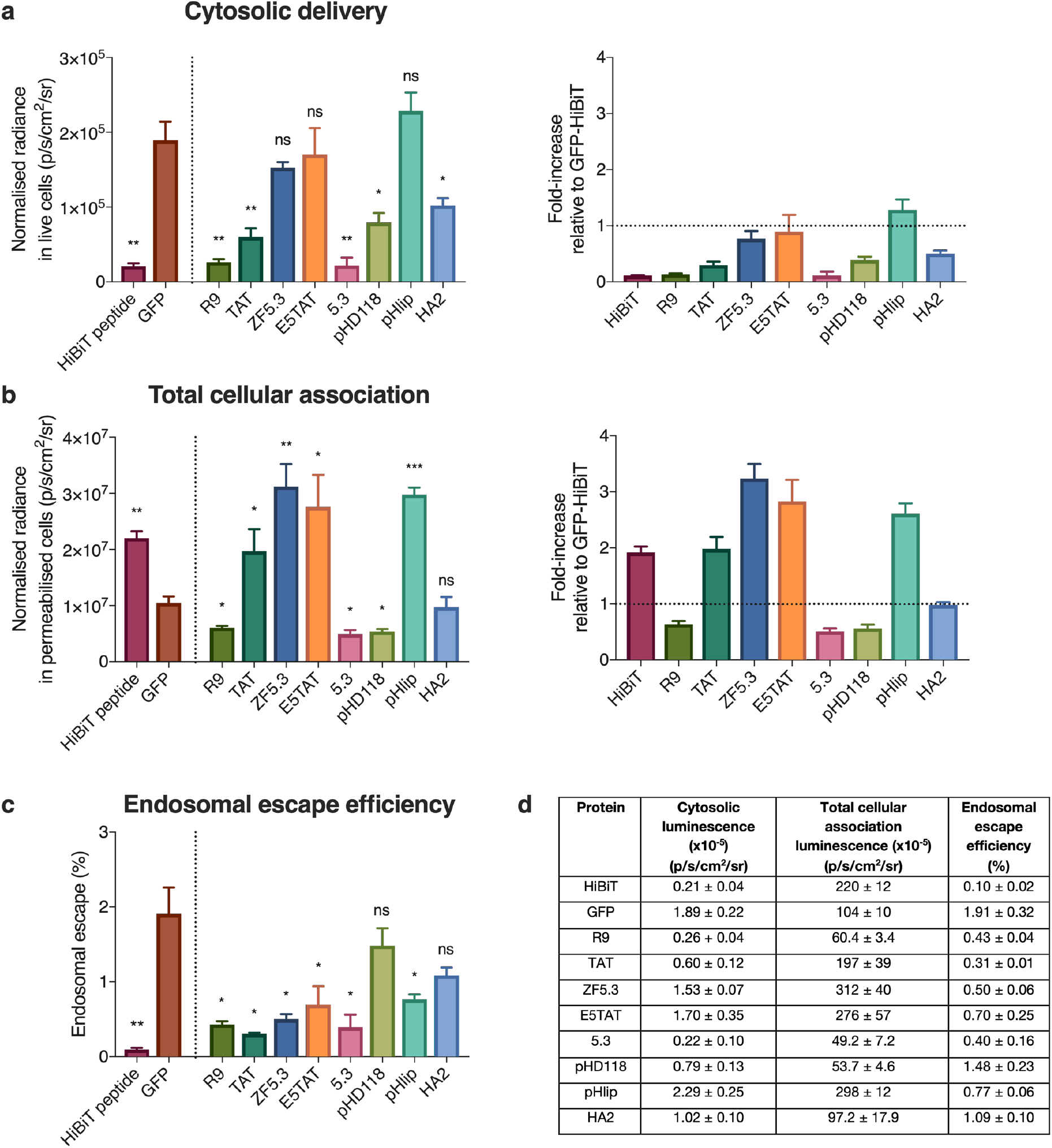
EEPs do not increase endosomal escape efficiency of GFP. a) Cytosolic luminescent signal of EEP-GFP-HiBiT in HEK293-LSA cells and fold-increase in signal with respect to GFP (represented by dotted line = 1). HEK293-LSA cells were incubated with EEP-GFP-HiBiT proteins at 1 μM for 4 hours. b) Total cellular association of EEP-GFP-HiBiT in HEK293 cells and fold-increase with respect to GFP-HiBiT (represented by dotted line = 1) determined by permeabilising the cells using 0.01% w/v digitonin. c) Endosomal escape efficiency of EEP-GFP-HiBiT proteins determined by ratioing cytosolic signal with total cellular association. d) Summary of cytosolic luminescence, total cellular association luminescence and endosomal escape efficiency for all proteins. Student's t test was used to compare differences between EEP-GFP-HiBiT fusion protein and GFP. Data represents mean ± SEM, n=3. ns (not significant) denotes p ≤ 0.05, * denotes p ≤ 0.05, ** denotes p ≤ 0.01 and *** denotes p ≤ 0.001.

When the concentration of protein was adjusted to give similar total cellular association, the positively charged EEPs no longer exhibited increased cytosolic accumulation. In fact, these EEPs exhibited the same or significantly less cytosolic delivery than GFP. This translates to a significantly lower percentage of endosomal escape compared to GFP without EEP (Fig. 6c and d), which is consistent with the endosomal escape percentages shown in Fig. 4d when cells were treated with equal concentrations of GFP. HiBiT peptide alone showed the lowest endosomal escape efficiency (Fig. 6c and d), indicating that HiBiT does not possess endosomal escape properties. Similar results were observed for HeLa cells (Supplementary Fig. 7). Taken together, these results suggest that EEPs do not improve endosomal escape efficiency, but rather just increase cell association.

## Discussion

The process of endosomal escape is postulated to be highly inefficient, however until now, our estimates of endosomal escape efficiency have been qualitative. Image based techniques to measure endosomal escape are inherently subjective and qualitative. The majority of material delivered to the cell is concentrated within endosomes and lysosome, as highlighted by the punctate fluorescence observed in Fig. 5. The signal from these compartments can easily swamp the diffuse signal from material that has escaped from the endosome. Furthermore, the signal from out of focus endosomal compartments is often confused with endosomal escape. The images in Figs 5 and S5 highlight the limitations of using fluorescence microscopy to investigate endosomal escape. It is challenging to compare images with such highly contrasting intensities, and it is also challenging to observe weak cytosolic signal, as the bright punctate fluorescence in the endosomes overwhelms any diffuse cytosolic signal.

To quantify endosomal escape, sensors that signal when cargo reaches the cytosol are required. The first generation of ‘switch on’ endosomal escape sensors were based on a split GFP^16,17,19^. This sensor significantly improves the ability to detect endosomal escape. However, as with fluorescence microscopy, the limited sensitivity of fluorescence techniques means the limit of detection is high and concentration of cargo delivered to the cells needs to be much higher than the therapeutically relevant doses. This means that endosomal escape within a therapeutically relevant window cannot be studied. A much more sensitive probe based on inactivated Renilla luciferase (RLuc) protein that is activated upon deglycosylation in the cytosol has recently been reported^33,34^. This method enables correlation of endosomal escape with transfection efficiency of the delivered mRNA, but does not quantify endosomal escape efficiency, and employs a large glycosylated protein (~75 kDa) as the sensor. Here, we have demonstrated that SLEEQ is a robust and highly sensitive assay that allows direct quantification of endosomal escape using a short peptide sensor. The assay is greater than 4 orders of magnitude more sensitive than a split GFP system (Fig. 2). The limit of detection for the SLEEQ assay is 2.1 x 10^4^ p/s/cm^2^/sr, which is > 8 times lower that the signal detected for GFP (GFP = 1.89 x 10^5^ p/s/cm^2^/sr).

Our results support the contention that endosomal escape is an inefficient process. Only ~2% of GFP associated with HEK293 cells was able to gain entry into the cytosol. This demonstrates not only the inefficiency of endosomal escape, but importantly highlights the sensitivity of SLEEQ to be able to quantify such low levels of endosomal escape. We also investigated endosomal escape in HeLa cells, and found they exhibited significantly higher endogenous levels of endosomal escape (~7%) than the HEK293 cells. This suggests that HeLa cells may have naturally leakier endosomes than HEK293 cells. These results demonstrate differences in intracellular trafficking in different cell lines and highlights the importance of studying different cell types.

The most notable result from the SLEEQ assay is that none of the putative EEPs increased the endosomal escape efficiency of the protein they were fused to. When dosed at 1 μM, positively charged peptides (R9, TAT, ZF5.3, and E5TAT) enhanced cytosolic delivery of GFP-HiBiT in both HEK293 (7 to 30-fold increase) and HeLa (7 to 21-fold increase) cells (Fig. 4a and Supplementary Fig. 5a). Although 5.3 peptide is also positively charged, it did not appear to significantly accumulate in the cytosol of HEK293 cells, but showed significant cytosolic delivery in HeLa cells. The positively charged peptides increased association with cells by an even greater amount (Fig. 4b and Supplementary Fig. 5b). When the concentration of the proteins was lowered to match the association of GFP, the positively charged peptides showed significantly lower cytosolic delivery than unmodified GFP. This demonstrates two important points. 1) The increased cytosolic delivery of cationic EEPs seen at 1 μM concentration (Fig. 4a) can be attributed to increase accumulation with the cells, likely driven by electrostatic interactions with the negatively charged plasma membrane. There is no evidence to suggest that these cationic peptides play an active role in inducing endosomal escape. 2) The positive charge of the EEPs actually hinders the cytosolic delivery when similar amounts of protein are associated with the cells (Fig. 6a). It is likely that the electrostatic interaction with the EEPs and the negatively charged membrane is retained in the endosome and they are unable to dissociate from the endosomal membranes, which inhibits the endogenous leakage of proteins into the cytosol. Negatively charged EEPs (pHD118, pHlip and HA2) showed no increase in cytosolic delivery and no significant increase in their total cellular association.

These results fundamentally change our understanding of the role of EEPs and how they promote cytosolic delivery. Rather than inducing an active endosomal escape mechanism, it appears that when positively charged EEPs are fused to a protein, they increase cytosolic delivery simply by increasing cellular accumulation. The increased cytosolic delivery comes at the cost of reduced endosomal escape efficiency, as the majority of protein appears to remain electrostatically bound to the membrane.

In summary, SLEEQ is a highly sensitive, direct and quantitative method for detecting endosomal escape. We demonstrated that endosomal escape is an inefficient process that varies between cell lines. In an effort to enhance endosomal escape, we tested a range of EEPs that have been widely studied to enhance cytosolic delivery. Our findings demonstrate that while positive EEPs improved cytosolic accumulation, it was not via an active mechanism of transport. Increased cytosolic accumulation was a result of an increase in total association of EEPs with cells. For the first time, SLEEQ enables the detection of constitutive levels of cytosolic transport and is a powerful assay that has the potential to quantify endosomal escape of a range of biological therapies at therapeutically relevant concentrations.

## Supporting information

Supplementary information, including protein characterisation and assay validation is available for download.

## Acknowledgements

MALDI was performed at the Melbourne Centre for Nanofabrication (MCN) in the Victorian Node of the Australian National Fabrication Facility (ANFF). We thank Dr David Rudd for providing his MALDI expertise.

## Author Contributions

C.W.P. and A.P.R.J. developed the concept and supervised the study. S.L.Y.T. performed fluorescence imaging for visualising actin filaments, designed and performed all of the luminescence experiments, analysed the data and designed the Fig.s. J.J.R. assisted with cloning, virus production, prepared and characterised the fusion proteins and performed the imaging for visualisation protein distribution. H. A. W. provided assistance in virus production and developing the concept. D.Y. provided assistance in designing molecular constructs of the fusion proteins. S.L.Y.T. wrote the manuscript, J.J.R., D.Y., C.W.P. and A.P.R.J. edited the manuscript.

## Methods

### Plasmid construction

All plasmids were constructed using NEBuilder HiFi DNA assembly master mix (NEB) with PCR products, vector restriction digests or DNA oligonucleotides with compatible overhangs. Cloning was performed in TOP10 chemically competent *Escherichia coli* (E. coli) (Thermo Fisher Scientific).

GFP (muGFP)^35^ with a C-terminal HiBit peptide (NlucC variant #86: VSGWRLFKKIS)^18^ was fused to the C-terminus of 14 x His bdSUMO^36^ and inserted into pET His6 MBP TEV LIC cloning vector (2M-T), a gift from Scott Gradia (RRID:Addgene_29708). The vector’s 6 x His MBP TEV coding sequence was replaced with 14 x His bdSUMO – GFP – HiBiT. Endosomal escape peptide sequences were purchased as DNA oligonucleotides from Integrated DNA Technologies (IDT) and inserted between bdSUMO and GFP. An additional glycine was inserted before R9 to improve bdSUMO cleavage.

pSF1389 encoding bdSENP1 was a gift from Dirk Görlich (RRID:Addgene_104962). The sequence encoding GFP_1-10_ for the split-GFP assay^37^ was ordered as a gBlock from IDT and was also inserted into pET His6 MBP TEV LIC cloning vector (2M-T) where the MBP sequence was replaced with GFP_1-10_.

The LgBiT-SNAP-actin lentiviral plasmid was constructed by insertion of LgBiT DNA^18^, SNAP-tag DNA (pSNAPf, New England Biolabs) and β-Actin DNA (Actin mRFP-PAGFP was a gift from Guillaume Charras & Tim Mitchison, RRID:Addgene_62382) into the third-generation lentiviral plasmid pCDH-EF1-IRES-Puro (System Biosciences).

### Cell culture

Dulbecco’s Modified Eagle Medium (DMEM) (4.5g/L glucose, 110 mg/L sodium pyruvate, no glutamine), 100x GlutaMAX supplement, Dulbecco’s Phosphate-buffered saline (DPBS, no calcium, no magnesium) and TrypLE were purchased from Thermo Fisher Scientific.

Human embryonic kidney cells (HEK293, ATCC Cat# CRL-1573, RRID:CVCL_0045) and human cervical cancer epithelial cells (HeLa, a gift from David Jans) were cultured in DMEM supplemented with 10% fetal bovine serum (FBS), 1x GlutaMAX and 100 units/mL streptomycin and 100 μg/mL penicillin. Both cell lines were maintained at 37°C in a 5% CO_2_ atmosphere, and sub-cultured every 2-4 days when 70-90% confluency was reached.

HEK293 and HeLa cells that stably expresses LgBiT-SNAP-actin were maintained in DMEM supplemented with 10% FBS, 1x GlutaMAX and 100 units/mL streptomycin, 100 μg/mL penicillin and 2 μg/mL puromycin every 5 passages. Only cells with less than 30 passages were used in all experiments. Cells were tested negative for mycoplasma contamination.

### Generation of stable cell lines expressing LgBiT-SNAP-actin

Lentivirus was produced by transfecting HEK293-FT cells (Thermo Fisher Scientific) with the third-generation lentiviral vector system using lipofectamine 3000 (Thermo Fisher Scientific). Lentivirus was harvested 48 hours after transfection and applied to relevant cell lines. Cells transduced with lentivirus were grown until confluent then selected with 2 μg/mL puromycin for positive incorporation of the transfer gene. Resistant cells were then single cell sorted into 96-well plates using a MoFlo Astrios (Beckman Coulter) to begin clonal cell lines.

### Sensitivity of split NanoLuciferase and split GFP

The sensitivity of split NanoLuciferase and split GFP were assessed by combining a fixed amount of purified LgBiT or GFP_1-10_ protein with increasing molar equivalents of HiBiT or GFP_11_ peptide. For split NanoLuciferase, 25 uL of desired concentrations (half-log increases) of HiBiT peptide were combined with 50 μL of 100 nM LgBiT protein in a black 96-well clear bottom microplate. To the mixture, 25 μL of NanoGlo Live Cell Substrate (Promega) was added and luminescence was measured on the In Vivo Imaging System Lumina (IVIS) Lumina II (Perkin Elmer) 10 minutes after substrate addition. Luminescence data was processed with Living Image 4.3.1 software. Luminescence was quantified in average radiance in units of photons per second per centimetre squared per steradian (p/s/cm^2^/sr).

For split GFP, 50 μL of desired concentrations of GFP_11_ peptide (GL BioChem) were combined with 50 μL of 6 μM GFP_1-10_ for a final volume of 100 μL in a black 96-well clear bottom microplate. Incubation proceeded at 37°C inside ClarioSTAR microplate reader (BMG Labtech) with fluorescence emission at 515 nm being detected after 470 nm excitation 2.75 hours post GFP_11_ addition (the time at which peak fluorescence intensity was observed).

Radiance and fluorescence values recorded were averages of three experiments subtracted by the average radiance or fluorescence values of blank media or PBS.

### Digitonin permeabilisation

To determine the concentration of digitonin required for total cell permeabilization, HEK cells expressing LgBiT-SNAP-actin were seeded one day prior at 10,000 cells/well. Cells were incubated with HiBiT peptide at 1 μM for 2 hours. Cells were then washed thrice with cell growth media (DMEM supplemented with 10% FBS). Digitonin stock solution (prepared in DMSO as 10% w/v) was diluted and added to cells for final concentrations of 0.001, 0.005, 0.01, 0.02 and 0.05% w/v for 30 minutes at 37°C. NanoGlo Live Cell Substrate, prepared according to manufacturer’s instructions, was added to cells and luminescence was measured on the IVIS 30 minutes after substrate addition. Exposure time was set to 10 seconds.

### Secretion of LgBiT

To determine LgBiT secretion, cells expressing LgBiT only or LSA were seeded at 10,000 cells/well to a final volume of 100 μL in black 96-well clear bottom microplate and incubated at 37°C for 24 hours. 50 μL of the cell supernatant was removed carefully without touching the cells adhered to the bottom of the plate and transferred to another black 96-well clear bottom microplate. The remaining volume of media was discarded and the cells were washed once with media.

50 μL of 0.02% w/v digitonin and 25 μL of 4 nM HiBiT peptide (diluted in DMEM supplemented with 10% FBS) was added to cells. 25 μL of 4 nM HiBiT peptide was added to the collected cell supernatant. Both cells and cell media were incubated for 1 hour at 37°C. 25 uL of NanoGlo Live Cell Substrate was added to both cell media and cells and luminescence was measured on IVIS Lumina II 10 minutes after substrate addition. Exposure time was set to 10 seconds. Cells and cell supernatant that were not treated with HiBiT peptide were blank samples. Percentage of LgBiT secretion from cells was calculated according to the equation below.

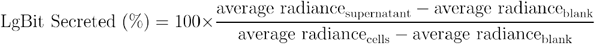

### Calcein uptake

Wild type HEK293 and HEK293-LSA cells were seeded at 40,000 cells/well in a 96-well plate 1 day prior to the experiment. The cells were treated with calcein (100 μg/mL) for 0.5, 1, and 2 hours. The cells were washed twice in cold growth media, then once with cold PBS, and detached with 50 μL TrypLE for 10 min. 100 μL 1% bovine serum albumin (BSA) in PBS was added to each well and the entire contents were transferred to a 96-well, V-bottom plate. The cells were spun at 400 g for 5 minutes and the supernatant discarded. The cell pellet was resuspended in 100 μL 1% BSA in PBS and analysed by flow cytometry (Stratedigm). Data processing was performed on FlowJo 10.

### Protein expression

Plasmids were freshly transformed into BL21 Star (DE3) *E. coli* (Thermo Fisher Scientific) prior to each expression batch. Transformed bacteria were directly inoculated into 2 L plastic baffled flasks (Thomson Instrument Company) containing 200 mL optimised growth medium with 15 g/L tryptone (Thermo Fisher Scientific), 30 g/L yeast extract (Thermo Fisher Scientific), 8 mL/L glycerol (Promega) and 10 g/L NaCl and shaken at 200 RPM overnight at 37°C. High density cultures were then reduced to room temperature and induced with 0.4 mM IPTG (Roche) for six hours. Bacteria were harvested by centrifugation at 4,000 *g*. Bacterial pellets were stored at −20°C if processing was not immediate. The full sequences of each protein can be found in Supplementary Fig. 8.

### Protein purification

Bacterial pellets were resuspended in a high salt buffer (1 M NaCl, 50 mM Imidazole, 50 mM monosodium phosphate, adjusted to pH 8.0) supplemented with complete EDTA free protease inhibitors, 2mM MgCl_2_ and benzonase. Resuspended bacteria were lysed by homogenisation with an EmulsiFlex-C3 (Avestin) before centrifugation at 12,000 *g* and clarified through a 0.45 μm syringe filter to remove cellular debris. Protein was purified by immobilised metal affinity chromatography (IMAC) using Protino Ni-NTA agarose (Machery-Nagel). Captured protein was washed copiously with high salt buffer and a low salt buffer (100 mM NaCl, 50 mM Imidazole, 50 mM monosodium phosphate, adjusted to pH 8.0) before elution (300 mM NaCl, 450 mM Imidazole, 50 mM monosodium phosphate, adjusted to pH 8.0). Eluted GFP fusion proteins were concentrated and buffer exchanged into buffer PB (500 mM NaCl, 50 mM Tris-HCl, pH 8.0) using 30 kDa Amicon centrifugal filters (Merck). Eluted bdSENP1, LgBiT and GFP_1-10_ were concentrated and buffer exchanged into PBS using a 10 kDa Amicon. Non-fluorescent protein concentration was quantified by A280 using estimated extinction coefficients. GFP fusion protein concentration was quantified by A490 using ε = 117,000 M^-1^cm^-1^^35^. bdSUMO cleavage from GFP proteins was achieved by a one hour 37°C incubation with bdSENP1 in a molar ratio of 300:1 in cleavage buffer (500mM NaCl, 50mM Tris-HCl, 250 mM sucrose, 2 mM MgSO_4_, 2 mM DTT, pH 8.0). bdSUMO, bdSENP1 and uncleaved GFP fusion proteins were separated by IMAC. Cleaved GFP proteins were buffer exchanged into buffer PB and concentrated using an Amicon 10 kDa MWCO ultrafiltration device. Cleaved ZF5.3 was buffer exchanged into buffer PB supplemented with 100 μM ZnCl_2_. Purity of the proteins were determined by performing SDS-PAGE under non-reducing conditions and MALDI mass spectrometry. All proteins were filtered using a 0.22 μm syringe filter then aliquoted and snap frozen in liquid nitrogen and stored at −80°C.

### Fluorescence microscopy

To visualise actin staining, HEK293-LSA cells were seeded in 8-well microscopy chambers (Ibidi) at 30,000 cells/well. After an overnight incubation, cells were treated with 1 μM SNAP-cell 647-SiR (New England Biolabs) for 30 minutes, then washed three times with fresh media and incubated for another 30 minutes to remove excess substrate. Cells were fixed using 4% paraformaldehyde for 10 minutes and washed thrice with PBS. Hoechst 33342 (Thermo Fisher Scientific) was diluted 1:2000 in FluoroBrite DMEM (Thermo Fisher Scientific) supplemented with 10% FBS and added to fixed cells. Imaging was performed using Olympus IX83 deconvolution wide field microscope with a 60x silicone objective. All images were processed using SlideBook 6.0 and ImageJ Fiji 2.0.0 software^38^.

For cell viability experiment (Supplementary Fig. 4), HEK293-LSA cells were seeded one day prior at 10,000 cells/well into black 96-well clear bottom plates. Cells were treated with 1 μM EEP-GFP-HiBiT proteins and free HiBiT peptide for 4 hours. Cells were washed with cell media three times and stained with Hoechst 33342 and propidium iodide (PI). Leica SP8 Lightning confocal microscope was used to image cells (Hoescht – 405 nm excitation, 420-470 nm emission, PI – 488 nm excitation, 500-600 nm emission).

To visualise cellular distribution of EEP-GFP-HiBiT proteins, HEK293-LSA and HeLa-LSA cells were seeded one day prior at 10,000 cells/well into black 96-well clear bottom plates. EEP-GFP-HiBiT fusion proteins were prediluted to 1 μM in DMEM supplemented with 10% FBS and added to cells for 4 hours. Cells were then washed with cell media three times and stained with Hoechst for 10 minutes before finally being imaged live in phenol-red free 10% FBS DMEM using a Leica TCS SP8 confocal microscope (GFP – 488 nm excitation, 500-570 nm emission, Hoechst – 810 nm (two photon) excitation, 420-470 nm emission).

### MALDI mass spectrometry

MALDI-MS was performed using a MALDI-TOF/TOF ultrafleXtreme (Bruker-Daltonics) equipped with a 1 kHz laser or a MALDI-7090 (Shimadzu) equipped with a 2 kHz laser, both operated in linear positive-ion mode. A total of 5,000 shots were summed using a 100 μm laser diameter and a user optimised laser intensity. Samples were prepared using the matrix 3-(4-hydroxy-3,5-dimethoxyphenyl)prop-2-enoic acid (sinapinic acid) and dissolved in a mixture of TA30 (30% acetonitrile and 70% water with trifluoroacetic acid (0.1% v/v)). 1 μL of each protein was initially combined with 10 μL of matrix before spotting 2 μL of the sample onto a ground steel plate or AnchorChip target. External calibration was achieved with bovine serum albumin (BSA) using the average mass of the [M+H]^+^ m/z ~66.5 kDa and [M+2H]^2+^ m/z ~33.3 kDa (Supplementary Table 1). Acquired spectra were exported to the opensource software mMass for processing (Supplementary Table 1 and 2).

### Activity of EEP-GFP-HiBiT fusion proteins

To determine the activity of the EEP-GFP-HiBiT fusion proteins, 50 nM of purified LgBiT protein was combined with 0.5 nM EEP-GFP-HiBiT fusion proteins in a black 96-well clear bottom microplate. NanoGlo Live Cell substrate was added to each well and luminescence was measured on the IVIS at 10 minutes post-substrate addition. Luminescent signal from EEP-GFP-HiBiT proteins were normalised to the reference HiBiT peptide signal (Fig. 3b).

### SLEEQ assay for determining endosomal escape efficiency

HEK293 cells stably expressing LgBiT-SNAP-actin (LSA) constructs were seeded one day prior at 10,000 cells per well into black 96-well clear bottom microplates. EEP-GFP-HiBiT proteins were added to cells at a final concentration of 1 μM and incubated for 4 hours. For matched association experiments, the concentrations EEP-GFP-HiBiT proteins were adjusted accordingly (HiBiT: 2 nM, GFP: 1 μM, R9: 2 nM, TAT: 4 nM, ZF5.3: 2 nM, E5TAT: 2 nM, 5.3: 2 nM, pHD118: 500 nM, pHlip: 1 μM and HA2: 300 nM. The cells were washed 3 times with fresh media (DMEM, no phenol red, supplemented with 10% FBS) to remove excess protein that had not been internalised.

To measure cytosolic signal, 25 μL of diluted NanoGlo live cell substrate and 75 μL fresh media (with equal volume of DMSO used for digitonin permeabilisation) was added to cells. Luminescence measurements were made at 10 minutes after substrate addition on IVIS Lumina II. Exposure time was set to 10 seconds. Immediately after measuring cytosolic signal, 50 μL of cell supernatant was carefully transferred without disturbing the cell layer into another black 96-well clear bottom microplate and its luminescent signal measured.

To measure total cellular association, cells were treated with 0.01% w/v digitonin after the three washes and incubated for 1 hour at 37°C before substrate addition and luminescence measurement.

Both cytosolic and total cellular association signal were adjusted by subtracting the blank signal (untreated cells) and normalised to their respective EEP-GFP-HiBiT activities (Fig. 3b). In addition, the signal in cell supernatant was also subtracted from the cytosolic signal to account for any small amount of LgBiT/HiBiT complex released into the supernatant. The equations are shown below.

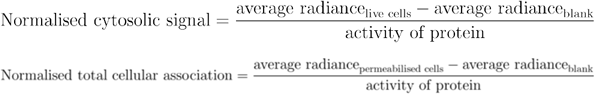

Endosomal escape efficiency was determined by the ratio of cytosolic signal to total cellular association. The equation is shown below.

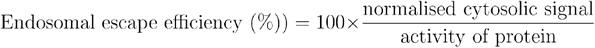

Endosomal escape efficiency was calculated for each independent experiment and averaged to provide mean ± SEM (Fig. 4c, d and Fig. 6c, d).

The limit of quantification was defined as three times the standard deviation of the background signal (LSA expressing cells in the presence of substrate, without HiBiT)

### Statistical analysis

Luminescence data was processed on GraphPad Prism 7 software and presented as mean ± standard error of mean (mean ± SEM). Statistical analyses were performed using unpaired, two-tailed Student’s t tests, with GFP being the control group. Sample sizes (n) are provided in the respective Fig. legends. Asterisks represent statistical significance (ns denotes p > 0.05, * denotes p ≤ 0.05, ** denotes p ≤ 0.01, *** denotes p ≤ 0.001 and **** denotes p ≤ 0.0001).

## References

1. Mitragotri, S., Burke, P. A. & Langer, R. Overcoming the challenges in administering biopharmaceuticals: formulation and delivery strategies. Nat. Rev. Drug Discov. 13, 655–672 (2014).

2. Torchilin, V. Intracellular delivery of protein and peptide therapeutics. Drug Discov. Today Technol. 5, (2009).

3. Shete, H. K., Prabhu, R. H. & Patravale, V. B. Endosomal Escape: A Bottleneck in Intracellular Delivery. J. Nanosci. Nanotechnol. 14, 460–474 (2014).

4. Dominska, M. & Dykxhoorn, D. M. Breaking down the barriers: siRNA delivery and endosome escape. J. Cell Sci. 123, 1183–1189 (2010).

5. Selby, L. I., Cortez-Jugo, C. M., Such, G. K. & Johnston, A. P. R. Nanoescapology: progress toward understanding the endosomal escape of polymeric nanoparticles. Wiley Interdiscip. Rev. Nanomedicine Nanobiotechnology 9, e1452 (2017).

6. Ye, J. et al. CPP-assisted intracellular drug delivery, what is next? Int. J. Mol. Sci. 17, 1–16 (2016).

7. Heitz, F., Morris, M. C. & Divita, G. Twenty years of cell - penetrating peptides: from molecular mechanisms to therapeutics. Br. J. Pharmacol. 157, 195–206 (2009).

8. Vivès, E., Brodin, P. & Lebleu, B. A Truncated HIV-1 Tat Protein Basic Domain Rapidly Translocates through the Plasma Membrane and Accumulates in the Cell Nucleus. J. Biol. Chem. 272, 16010–16017 (1997).

9. Richard, J. P. et al. Cell-penetrating peptides: A reevaluation of the mechanism of cellular uptake. J. Biol. Chem. 278, 585–590 (2003).

10. Salomone, F. et al. A novel chimeric cell-penetrating peptide with membrane-disruptive properties for efficient endosomal escape. J. Control. Release 163, 293–303 (2012).

11. Yang, S. T., Zaitseva, E., Chernomordik, L. V. & Melikov, K. Cell-penetrating peptide induces leaky fusion of liposomes containing late endosome-specific anionic lipid. Biophys. J. 99, 2525–2533 (2010).

12. Cesbron, Y., Shaheen, U., Free, P. & Lévy, R. TAT and HA2 facilitate cellular uptake of gold nanoparticles but do not lead to cytosolic localisation. PLoS One 10, 1–18 (2015).

13. Verdurmen, W. P. R., Mazlami, M. & Plückthun, A. A quantitative comparison of cytosolic delivery via different protein uptake systems. Sci. Rep. 7, 1–13 (2017).

14. Martens, T. F., Remaut, K., Demeester, J., De Smedt, S. C. & Braeckmans, K. Intracellular delivery of nanomaterials: How to catch endosomal escape in the act. Nano Today 9, 344–364 (2014).

15. Kim, J. S. et al. Quantitative assessment of cellular uptake and cytosolic access of antibody in living cells by an enhanced split GFP complementation assay. Biochem. Biophys. Res. Commun. 467, 771–777 (2015).

16. Milech, N. et al. GFP-complementation assay to detect functional CPP and protein delivery into living cells. Sci. Rep. 5, 18329 (2016).

17. Schmidt, S. et al. Detecting Cytosolic Peptide Delivery with the GFP Complementation Assay in the Low Micromolar Range. Angew. Chemie - Int. Ed. 54, 15105–15108 (2015).

18. Dixon, A. S. et al. NanoLuc Complementation Reporter Optimized for Accurate Measurement of Protein Interactions in Cells. ACS Chem. Biol. 11, 400–408 (2016).

19. Lönn, P. et al. Enhancing Endosomal Escape for Intracellular Delivery of Macromolecular Biologic Therapeutics. Sci. Rep. 6, 32301 (2016).

20. Frankel, A. D. & Pabo, C. O. Cellular uptake of the tat protein from human immunodeficiency virus. Cell 55, 1189–1193 (1988).

21. Green, M. & Loewenstein, P. M. Autonomous functional domains of chemically synthesized human immunodeficiency virus tat trans-activator protein. Cell 55, 1179–1188 (1988).

22. Mitchell, D. J., Steinman, L., Kim, D. T., Fathman, C. G. & Rothbard, J. B. Polyarginine enters cells more efficiently than other polycationic homopolymers. J. Pept. Res. 56, 318–325 (2000).

23. Najjar, K. et al. Unlocking Endosomal Entrapment with Supercharged Arginine-Rich Peptides. Bioconjug. Chem. 28, 2932–2941 (2017).

24. Appelbaum, J. S. et al. Arginine topology controls escape of minimally cationic proteins from early endosomes to the cytoplasm. Chem. Biol. 19, 819–830 (2012).

25. Wissner, R. F., Steinauer, A., Knox, S. L., Thompson, A. D. & Schepartz, A. Fluorescence Correlation Spectroscopy Reveals Efficient Cytosolic Delivery of Protein Cargo by Cell-Permeant Miniature Proteins. ACS Cent. Sci. 4, 1379–1393 (2018).

26. Hunt, J. F. et al. A Biophysical Study of Integral Membrane Protein Folding †. Biochemistry 36, 15156–15176 (1997).

27. Qiu, L. et al. Endolysosomal-Escape Nanovaccines through Adjuvant-Induced Tumor Antigen Assembly for Enhanced Effector CD8+ T Cell Activation. Small 14, 1–11 (2018).

28. Wiedman, G., Kim, S. Y., Zapata-Mercado, E., Wimley, W. C. & Hristova, K. pH-Triggered, Macromolecule-Sized Poration of Lipid Bilayers by Synthetically Evolved Peptides. J. Am. Chem. Soc. 139, 937–945 (2017).

29. Wharton, S. A., Martin, S. R., Ruigrok, R. W. H., Skehel, J. J. & Wiley, D. C. Membrane Fusion by Peptide Analogues of Influenza Virus Haemagglutinin. J. Gen. Virol. 69, 1847–1857 (1988).

30. Plank, C., Oberhauser, B., Mechtler, K., Koch, C. & Wagner, E. The influence of endosome-disruptive peptides on gene transfer using synthetic virus-like gene transfer systems. J. Biol. Chem. 269, 12918–12924 (1994).

31. Lee, Y. J., Erazo-Oliveras, A. & Pellois, J. P. Delivery of macromolecules into live cells by simple co-incubation with a peptide. ChemBioChem 11, 325–330 (2010).

32. McLaughlin, S. The electrostatic properties of membranes. Annu. Rev. Biophys. Biophys. Chem. 18, 113–136 (1989).

33. Jiang, Y. et al. Quantitating Endosomal Escape of a Library of Polymers for mRNA Delivery. Nano Lett. 20, 1117–1123 (2020).

34. Lu, Q., Grotzke, J. E. & Cresswell, P. A novel probe to assess cytosolic entry of exogenous proteins. Nat. Commun. 9, 1–11 (2018).

35. Scott, D. J. et al. A Novel Ultra-Stable, Monomeric Green Fluorescent Protein for Direct Volumetric Imaging of Whole Organs Using CLARITY. Sci. Rep. 8, 1–15 (2018).

36. Frey, S. & Görlich, D. A new set of highly efficient, tag-cleaving proteases for purifying recombinant proteins. J. Chromatogr. A 1337, 95–105 (2014).

37. Cabantous, S. & Waldo, G. S. In vivo and in vitro protein solubility assays using split GFP. Nat. Methods 3, 845–854 (2006).

38. Schindelin, J. et al. Fiji: an open-source platform for biological-image analysis. Nat. Methods 9, 676–682 (2012).

