## Supplementary information, including protein characterisation and assay validation is available for download. for "Unravelling cytosolic delivery of endosomal escape peptides with a quantitative endosomal escape assay (SLEEQ)"

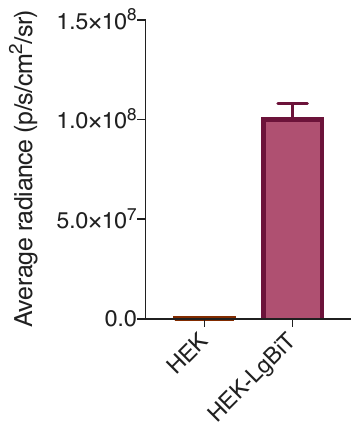

**Supplementary Fig. 1: Successful expression of LgBiT protein in HEK293 cells.** Non-transduced HEK293 cells and HEK293-LgBiT cells were seeded at 10,000 cells/well in black 96-well clear bottom plates. After overnight incubation, cells were treated with 1 nM HiBiT peptide and 0.01% wt/v digitonin for 1 hour at 37ºC to completely permeabilise the cells. NanoGlo Live Cell substrate was added to cells and luminescence was measured on IVIS Lumina II.

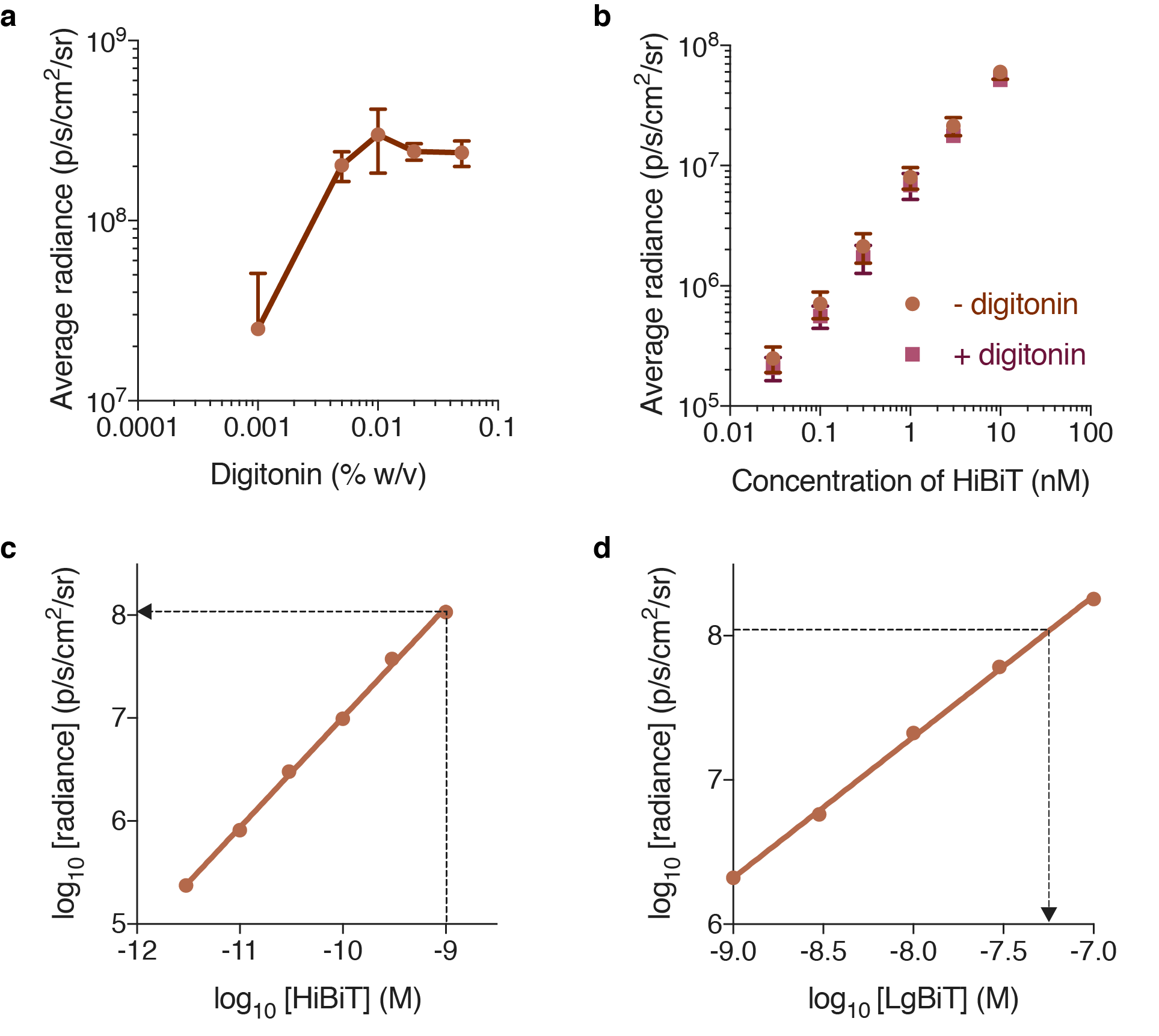

**Supplementary Fig. 2: Concentration of of LSA expressed in HEK293-LSA cells.** a) 0.01% wt/v digitonin is sufficient to achieve total cell permeabilization. HEK293-LSA cells were seeded at 10,000 cells/well in black 96-well plates. After overnight incubation, cells were incubated with 1 µM HiBiT peptide for 2 hours. Cells were washed and treated with various concentrations of digitonin (0.001, 0.005, 0.01, 0.02 and 0.05% w/v) for 30 minutes. NanoGlo Live Cell substrate was added and luminescence was measured on PerkinElmer In Vivo Imaging System Lumina II (IVIS) 30 minutes after substrate addition. Data represents mean ± SD, n =2. b) Digitonin does not affect luciferase activity. Purified LgBiT (50 nM) was combined with various concentrations of HiBiT peptide (0.03, 0.1, 0.3, 1, 3, 10 nM) with or without the presence of 0.01% wt/v digitonin. No difference in emitted radiance was observed with or without digitonin. Data represents mean ± SD, n=3. c) Linear luminescence in permeabilised HEK293-LSA cells was obtained over concentration range of 3 pM to 1 nM HiBiT. HEK293-LSA cells seeded in black 96-well clear bottom microplates at 10,000 cells/well were permeabilised with digitonin (0.01% wt/v) and treated with various concentrations of HiBiT peptide. Linear regression analysis was used to determine the line of best fit. Data represents mean ± SD, n=3. d) Total concentration of LSA expressed in HEK293 cells is determined to be 55.8 nM. Fixed 1 nM HiBiT peptide was titrated with various concentrations of purified LgBiT protein to construct a calibration curve. Data represents mean ± SD, n=2. The y-value at 1 nM HiBiT peptide in Fig. S2c (see arrow in Fig. S2c, y = 8.03) was interpolated to obtain the log_10_ [LgBiT] value in the curve obtained (see arrow in Fig. S2d, x = -7.25). The estimated concentration of LSA expressed in cells is therefore 10^-7.253^ = 55.8 nM.

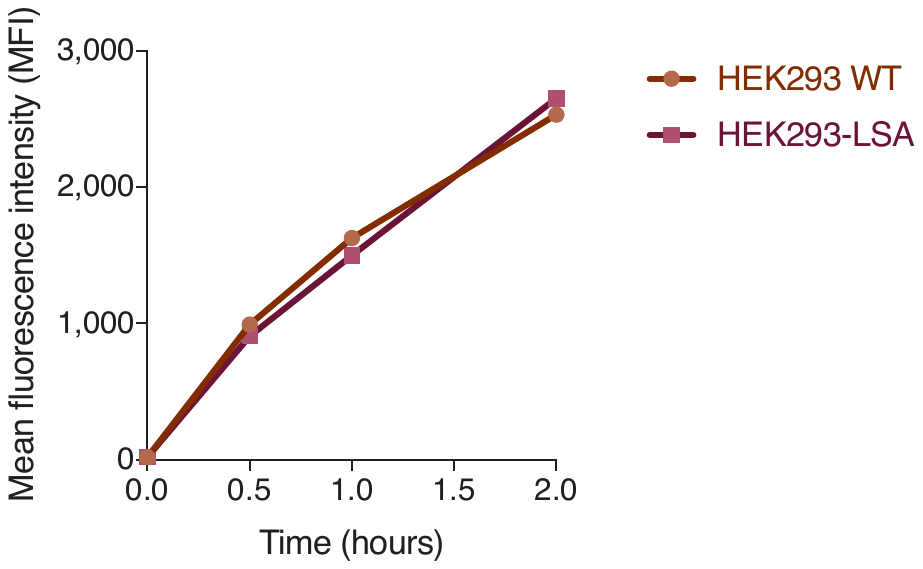

**Supplementary Fig. 3: Wild type (WT) HEK293 and HEK293-LSA cells show similar calcein uptake.**  HEK293 (WT) and HEK293-LSA cells were seeded at 40,000 cells/well in a 96-well plate. After overnight incubation, the cells were treated with calcein for 0.5, 1 and 2 hours respectively. The cells were washed and then analysed by flow cytometry.

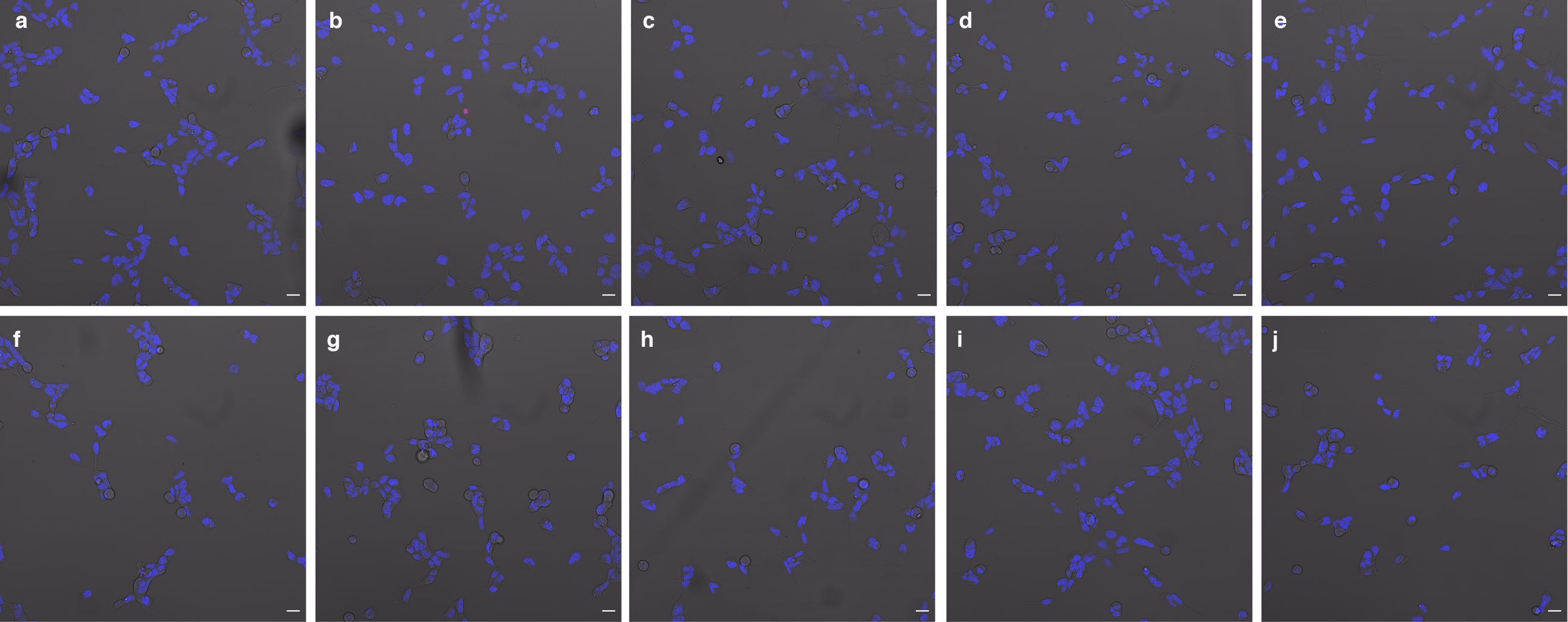

**Supplementary Fig. 4: HEK293-LSA cells maintain viability after treatment with EEP-GFP-HiBiT proteins.** After a 4-hour incubation with 1 µM EEP-GFP-HiBiT proteins or free HiBiT peptide: a) GFP, b) R9, c) TAT, d) ZF5.3, e) E5T**A**T, f) 5.3, g) pHD118, h) pHlip, i) HA2 and j) HiBiT, HEK293-LSA cells were washed and stained with Hoechst 33342 and propidium iodide (PI). Cells were imaged with confocal microscope. Hoechst 33342 stains all cell nuclei (blue) and PI only stains nuclei of dead cells (red). Fluorescence images are overlaid on bright-field image. Cells displayed their typical morphology, and less than 1% of the cells were positive for PI. ­

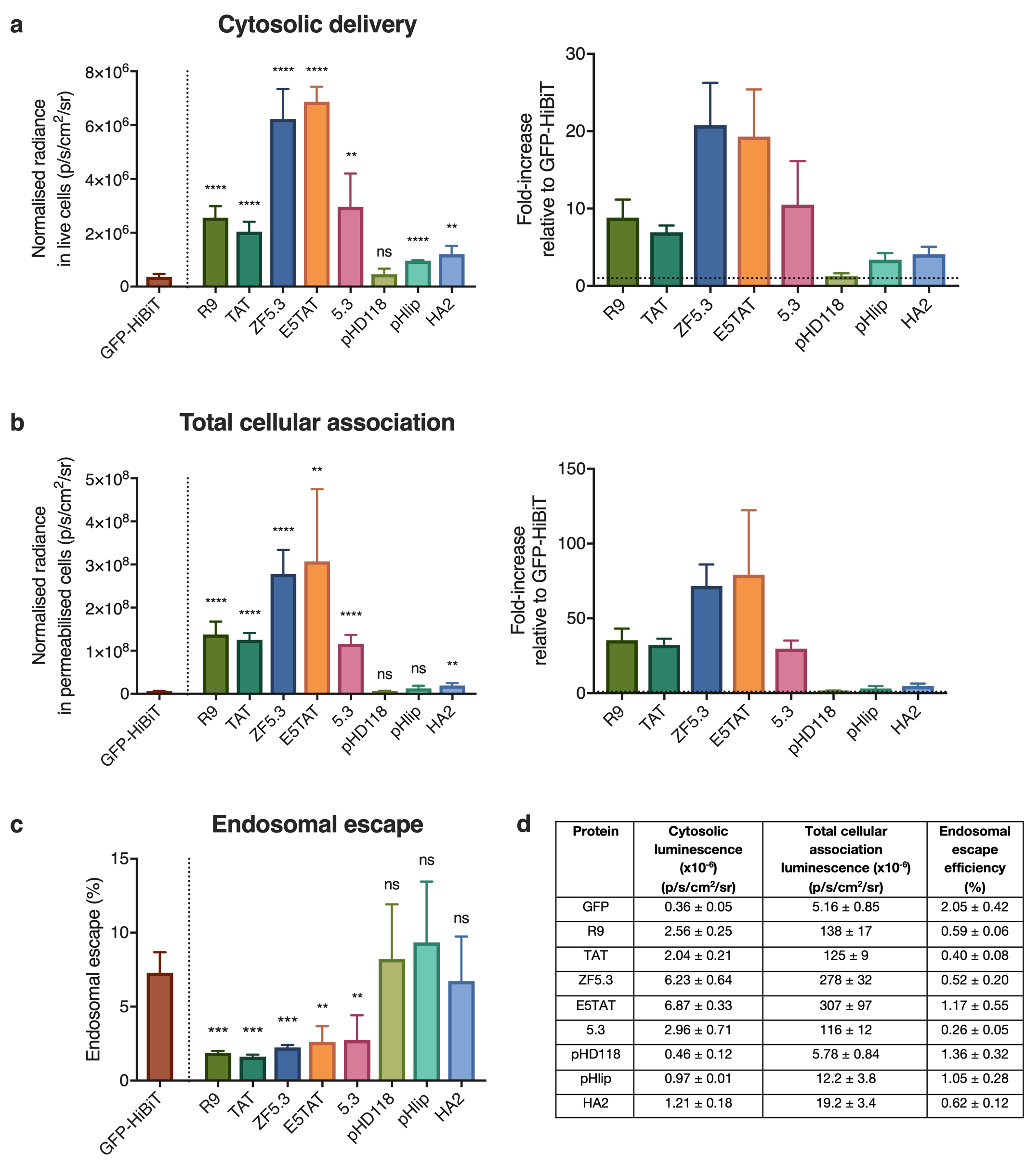

Supplementary Fig. 5: Cationic EEPs increase cytosolic delivery of GFP but do not increase endosomal escape efficiency in HeLa cells*.* a) Cytosolic luminescent signal of EEP-GFP-HiBiT in HeLa-LSA cells and fold-increase in signal with respect to GFP (represented by dotted line = 1). HeLa-LSA cells were incubated with EEP-GFP-HiBiT proteins at 1 µM for 4 hours. b) Total cellular association of EEP-GFP-HiBiT in HeLa cells and fold-increase with respect to GFP (represented by dotted line = 1) determined by permeabilising the cells using 0.01% wt/v digitonin. c) Endosomal escape efficiency of EEP-GFP-HiBiT proteins determined by ratioing cytosolic signal with total cellular association. d) Summary of cytosolic luminescence, total cellular association luminescence and endosomal escape efficiency for all proteins. Data represents mean ± SEM. Student’s t test was used to compare differences between EEP-GFP-HiBiT and GFP. n=3. ns denote p > 0.05, * denotes p < 0.05, ** denotes p < 0.01 and *** denotes p < 0.001

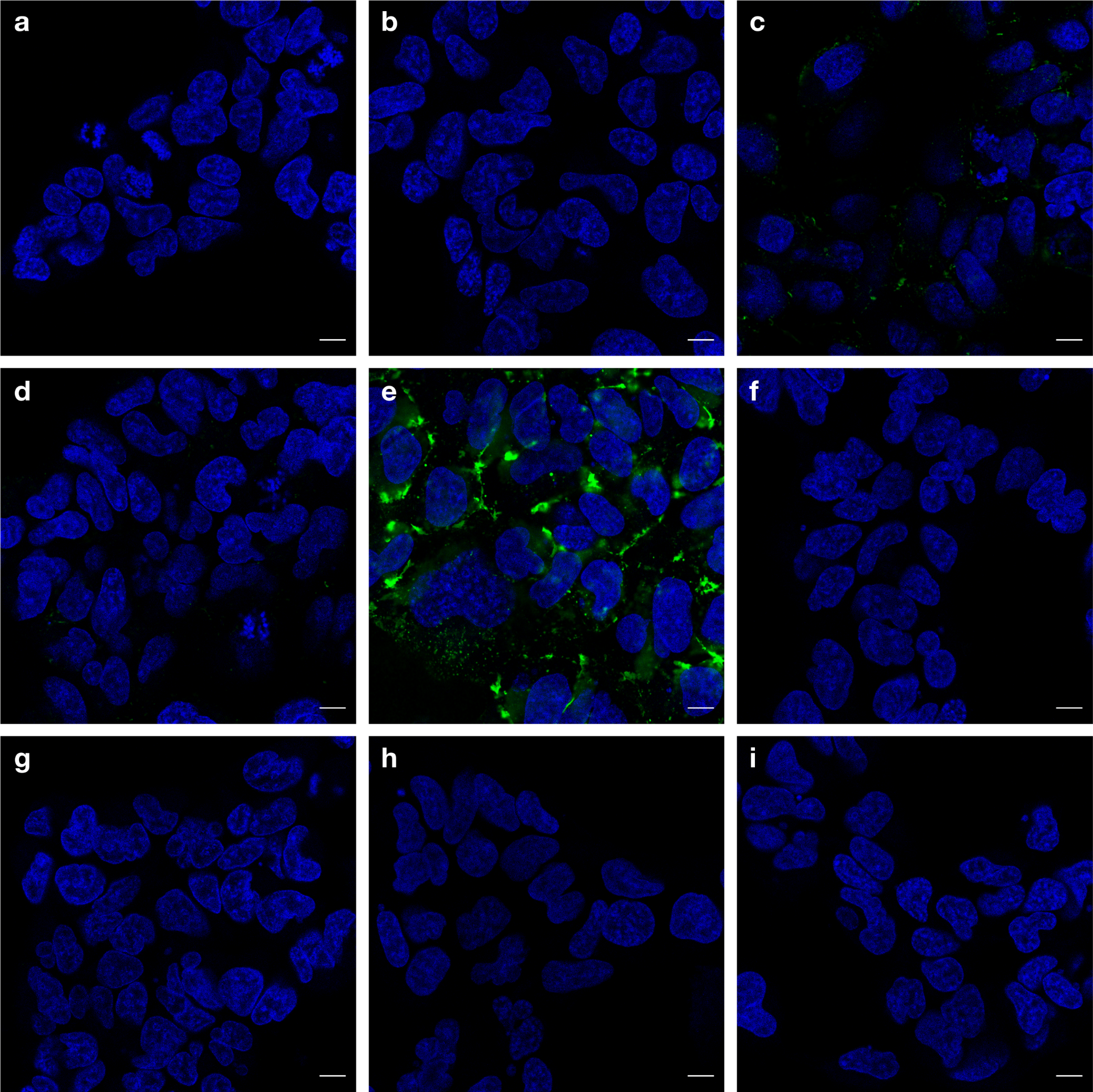

Supplementary Fig. 6: EEP-GFP-HiBiT fusion proteins exhibit punctate staining, suggesting limited endosomal escape. HEK293-LSA cells treated with EEP-GFP-HiBiT (green) proteins: a) GFP, b) R9, c) TAT, d) ZF5.3, e) E5TAT, f) 5.3, g) pHD118, h) pHlip and i) HA2 at 1 µM for 4 hours. Cells were washed and nuclei was stained with Hoechst 33342 (blue) before confocal microscopy. Images displayed with the same dynamic range to enable comparison of intensity. Scale bar = 10 µm

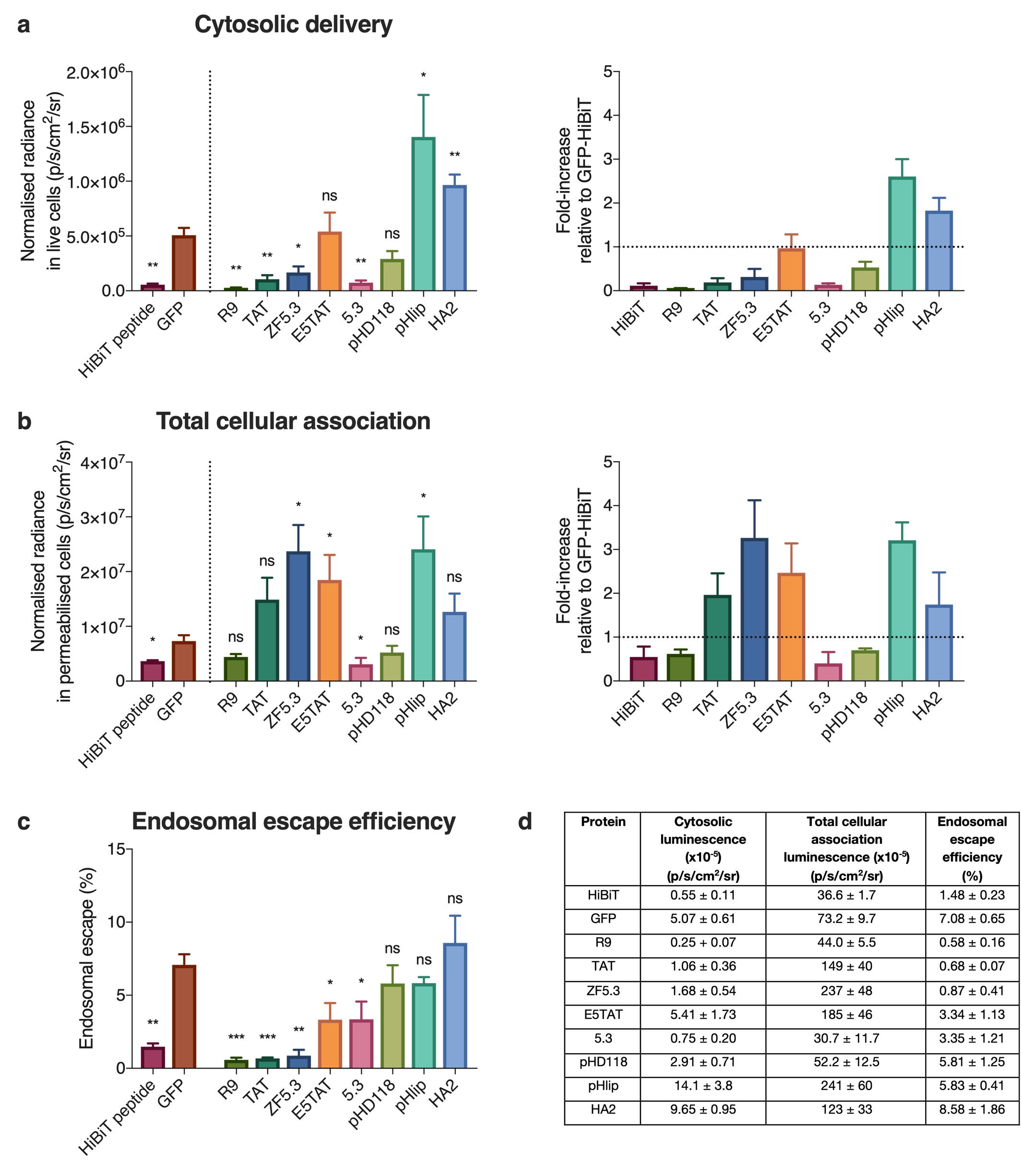

Supplementary Fig. 7: EEPs do not increase endosomal escape efficiency of GFP in HeLa cells*.* a) Cytosolic luminescent signal of EEP-GFP-HiBiT in HeLa-LSA cells and fold-increase in signal with respect to GFP (represented by dotted line = 1). HeLa-LSA cells were incubated with EEP-GFP-HiBiT proteins at 1 µM for 4 hours. b) Total cellular association of EEP-GFP-HiBiT in HeLa cells and fold-increase with respect to GFP (represented by dotted line = 1) determined by permeabilising the cells using 0.01% wt/v digitonin. c) Endosomal escape efficiency of EEP-GFP-HiBiT proteins determined by ratioing cytosolic signal with total cellular association. d) Summary of cytosolic luminescence, total cellular association luminescence and endosomal escape efficiency for all proteins. Data represents mean ± SEM. Student’s t test was used to compare the differences between EEP-GFP-HiBiT and GFP. n=3. ns denote p > 0.05, * denotes p < 0.05, ** denotes p < 0.01 and *** denotes p < 0.001

GFP

MSKHHHHSGHHHTGHHHHSGSHHHTGHINLKVKGQDGNEVFFRIKRSTQLKKLMNAYCDRQSVDMTAIAFLFDGRRLRAEQTPDELEMEDGDEIDAMLHQTGGSKGEELFTGVVPILVELDGDVNGHKFSVRGEGEGDATNGKLTLKFICTTGKLPVPWPTLVTTLTYGVLCFSRYPDHMKRHDFFKSAMPEGYVQERTISFKDDGTYKTRAEVKFEGDTLVNRIELKGIDFKEDGNILGHKLEYNFNSHNVYITADKQKNGIKAYFKIRHNVEDGSVQLADHYQQNTPIGDGPVLLPDNHYLSTQSVLSKDPNEKRDHMVLLEDVTAAGITHGMDELYKVSGWRLFKKISGGSG

R9

MSKHHHHSGHHHTGHHHHSGSHHHTGHINLKVKGQDGNEVFFRIKRSTQLKKLMNAYCDRQSVDMTAIAFLFDGRRLRAEQTPDELEMEDGDEIDAMLHQTGGGRRRRRRRRRSKGEELFTGVVPILVELDGDVNGHKFSVRGEGEGDATNGKLTLKFICTTGKLPVPWPTLVTTLTYGVLCFSRYPDHMKRHDFFKSAMPEGYVQERTISFKDDGTYKTRAEVKFEGDTLVNRIELKGIDFKEDGNILGHKLEYNFNSHNVYITADKQKNGIKAYFKIRHNVEDGSVQLADHYQQNTPIGDGPVLLPDNHYLSTQSVLSKDPNEKRDHMVLLEDVTAAGITHGMDELYKVSGWRLFKKISGGSG

TAT

MSKHHHHSGHHHTGHHHHSGSHHHTGHINLKVKGQDGNEVFFRIKRSTQLKKLMNAYCDRQSVDMTAIAFLFDGRRLRAEQTPDELEMEDGDEIDAMLHQTGGGRKKRRQRRRPPQASSKGEELFTGVVPILVELDGDVNGHKFSVRGEGEGDATNGKLTLKFICTTGKLPVPWPTLVTTLTYGVLCFSRYPDHMKRHDFFKSAMPEGYVQERTISFKDDGTYKTRAEVKFEGDTLVNRIELKGIDFKEDGNILGHKLEYNFNSHNVYITADKQKNGIKAYFKIRHNVEDGSVQLADHYQQNTPIGDGPVLLPDNHYLSTQSVLSKDPNEKRDHMVLLEDVTAAGITHGMDELYKVSGWRLFKKISGGSG

ZF5.3

MSKHHHHSGHHHTGHHHHSGSHHHTGHINLKVKGQDGNEVFFRIKRSTQLKKLMNAYCDRQSVDMTAIAFLFDGRRLRAEQTPDELEMEDGDEIDAMLHQTGGWYSCNVCGKAFVLSRHLNRHLRVHRRATASSKGEELFTGVVPILVELDGDVNGHKFSVRGEGEGDATNGKLTLKFICTTGKLPVPWPTLVTTLTYGVLCFSRYPDHMKRHDFFKSAMPEGYVQERTISFKDDGTYKTRAEVKFEGDTLVNRIELKGIDFKEDGNILGHKLEYNFNSHNVYITADKQKNGIKAYFKIRHNVEDGSVQLADHYQQNTPIGDGPVLLPDNHYLSTQSVLSKDPNEKRDHMVLLEDVTAAGITHGMDELYKVSGWRLFKKISGGSG

E5TAT

MSKHHHHSGHHHTGHHHHSGSHHHTGHINLKVKGQDGNEVFFRIKRSTQLKKLMNAYCDRQSVDMTAIAFLFDGRRLRAEQTPDELEMEDGDEIDAMLHQTGGGLFEAIAEFIENGWEGLIEGMGRKKRRQRRRPPQASSKGEELFTGVVPILVELDGDVNGHKFSVRGEGEGDATNGKLTLKFICTTGKLPVPWPTLVTTLTYGVLCFSRYPDHMKRHDFFKSAMPEGYVQERTISFKDDGTYKTRAEVKFEGDTLVNRIELKGIDFKEDGNILGHKLEYNFNSHNVYITADKQKNGIKAYFKIRHNVEDGSVQLADHYQQNTPIGDGPVLLPDNHYLSTQSVLSKDPNEKRDHMVLLEDVTAAGITHGMDELYKVSGWRLFKKISGGSG

5.3

MSKHHHHSGHHHTGHHHHSGSHHHTGHINLKVKGQDGNEVFFRIKRSTQLKKLMNAYCDRQSVDMTAIAFLFDGRRLRAEQTPDELEMEDGDEIDAMLHQTGGGPSQPTYPGDDAPVRDLIRFYRDLRRYLNVVTRHRYASSKGEELFTGVVPILVELDGDVNGHKFSVRGEGEGDATNGKLTLKFICTTGKLPVPWPTLVTTLTYGVLCFSRYPDHMKRHDFFKSAMPEGYVQERTISFKDDGTYKTRAEVKFEGDTLVNRIELKGIDFKEDGNILGHKLEYNFNSHNVYITADKQKNGIKAYFKIRHNVEDGSVQLADHYQQNTPIGDGPVLLPDNHYLSTQSVLSKDPNEKRDHMVLLEDVTAAGITHGMDELYKVSGWRLFKKISGGSG

pHD118

MSKHHHHSGHHHTGHHHHSGSHHHTGHINLKVKGQDGNEVFFRIKRSTQLKKLMNAYCDRQSVDMTAIAFLFDGRRLRAEQTPDELEMEDGDEIDAMLHQTGGIGEVLHELADDLPELQSWIKAAQQLSKGEELFTGVVPILVELDGDVNGHKFSVRGEGEGDATNGKLTLKFICTTGKLPVPWPTLVTTLTYGVLCFSRYPDHMKRHDFFKSAMPEGYVQERTISFKDDGTYKTRAEVKFEGDTLVNRIELKGIDFKEDGNILGHKLEYNFNSHNVYITADKQKNGIKAYFKIRHNVEDGSVQLADHYQQNTPIGDGPVLLPDNHYLSTQSVLSKDPNEKRDHMVLLEDVTAAGITHGMDELYKVSGWRLFKKISGGSG

pHlip

MSKHHHHSGHHHTGHHHHSGSHHHTGHINLKVKGQDGNEVFFRIKRSTQLKKLMNAYCDRQSVDMTAIAFLFDGRRLRAEQTPDELEMEDGDEIDAMLHQTGGAEEQQPWAQYLELLFPTETLLLEWGSKGEELFTGVVPILVELDGDVNGHKFSVRGEGEGDATNGKLTLKFICTTGKLPVPWPTLVTTLTYGVLCFSRYPDHMKRHDFFKSAMPEGYVQERTISFKDDGTYKTRAEVKFEGDTLVNRIELKGIDFKEDGNILGHKLEYNFNSHNVYITADKQKNGIKAYFKIRHNVEDGSVQLADHYQQNTPIGDGPVLLPDNHYLSTQSVLSKDPNEKRDHMVLLEDVTAAGITHGMDELYKVSGWRLFKKISGGSG

HA2

MSKHHHHSGHHHTGHHHHSGSHHHTGHINLKVKGQDGNEVFFRIKRSTQLKKLMNAYCDRQSVDMTAIAFLFDGRRLRAEQTPDELEMEDGDEIDAMLHQTGGGLFEAIEGFIENGWEGMIDGWYGASSKGEELFTGVVPILVELDGDVNGHKFSVRGEGEGDATNGKLTLKFICTTGKLPVPWPTLVTTLTYGVLCFSRYPDHMKRHDFFKSAMPEGYVQERTISFKDDGTYKTRAEVKFEGDTLVNRIELKGIDFKEDGNILGHKLEYNFNSHNVYITADKQKNGIKAYFKIRHNVEDGSVQLADHYQQNTPIGDGPVLLPDNHYLSTQSVLSKDPNEKRDHMVLLEDVTAAGITHGMDELYKVSGWRLFKKISGGSG

**Supplementary Fig. 8: Full sequences of uncleaved protein samples used in this study.** Sequences are coded as following: grey = 14xHis purification tag and bdSUMO; blue and underlined = endosomal escape peptide sequence; green = muGFP; yellow = HiBiT; none = amino acids intended to protect HiBiT peptide from C-terminal degradation. Note: bdSENP1 cleaves after bdSUMO C-terminal GG.

| **BSA** | **Expected average mass (kDa)** | **Observed mass (M/Z)** |
| --- | --- | --- |
| **[M+H]^+^** | ~ 66.5 | 66777.4 |
| **[M+2H]^+2^** | ~ 33.3 | 33255.6 |

**Supplementary Table 1:** Calibration of the MALDI instrument was performed with BSA. Observed masses are peak centroid values, expected masses were observed within the peak which is relatively broad due to isotopic combinations.

|  | **GFP** | **R9** | **TAT** | **ZF5.3** | **E5TAT** | **5.3** | **pHD118** | **pHlip** | **HA2** |
| --- | --- | --- | --- | --- | --- | --- | --- | --- | --- |
| **Expected average mass [M+H]^+^ (Da)** | 28212.9 | 29675.6 | 30072 | 31733.9 | 32379.6 | 32716.9 | 31012 | 31200.2 | 30943.8 |
| **Observed mass (M/Z)** | **28075.8** | **29524.9** | **29936.2** | **31587.0** | **32227.7** | **32578.4** | **31370.6** | **31058.5** | **30811.9** |
| **Minor Fragments (corresponding amino acids lost from C-terminus)** |  |  | *29478.0*  *(-ISGGSG)* | *31135.6*  *(-ISGGSG)* |  | *31865.9*  *(-KKISGGSG)* |  | *30595.0*  *(-ISGGSG)* |  |
|  |  |  | *29222.2*  *(-KKISGGSG)* | *30869.5*  *(-KKISGGSG)* |  |  |  | *30467.7*  *(-KISGGSG)* |  |
|  |  |  |  | *30732.1*  *(-FKKISGGSG)* |  |  |  | *30347.1*  *(-KKISGGSG)* |  |
|  |  |  |  |  |  |  |  | *30195.0*  *(FKKISGGSG)* |  |
|  |  |  |  |  |  |  |  | *30084.4*  *(-LFKKISGGSG)* |  |

**Supplementary Table 2:** **MALDI mass spectroscopy peak analysis of EEP-GFP-HiBiT proteins.** Observed masses are peak centroid values. Expected masses were observed within the peak which is relatively broad due to isotopic combinations. This is consistent with the difference in mass observed for BSA (Supplementary Table 1). The unusually broad appearance of pHD118 results in a centroid peak value that is greater than all other proteins, however the expected average mass is found within this broad peak. The minor fragments observed for some proteins correlate with a sequential loss of amino acids from the C-terminus, which is expected to be laser induced. Any true truncation which could affect the activity of HiBiT is compensated for by measuring the relative activity of each protein (Fig. 3).

| **Peptide sequence** | **Mass of fragment [M+H]^+^** |
| --- | --- |
| LFKKISGGSG | 976.2 |
| FKKISGGSG | 863.0 |
| KKISGGSG | 715.8 |
| KISGGSG | 587.6 |
| ISGGSG | 459.5 |

**Supplementary Table 3:** Sequential peptide fragments from the C-terminus of EEP-GFP-HiBiT proteins which correlate with fragments observed in Supplementary Table 2.
